# Human internal exposures of bisphenol A and six data-poor analogues predicted by physiologically based kinetic models with multimodal parameterization

**DOI:** 10.1101/2024.11.20.624474

**Authors:** Hélène Bigonne, Amrei Rolof, Inga Potapova, Shana J. Sturla, Georg Aichinger

## Abstract

**Background:** Bisphenols (BP) AF, B, E, F, M, and S have been introduced as substitutes for bisphenol A (BPA) and are increasingly used in consumer products. Despite widespread human exposure and potential adverse health outcomes related to BPF, BPB, BPS, and BPAF, their physiological disposition in humans is poorly characterized, which hinders assessment of associated risks.

**Objectives:** Our goal was to simulate the kinetic behavior of prevalent bisphenol analogs in organs of toxicological interest. To enable predictions of physiologically relevant internal concentrations of a family of structurally similar compounds with limited available human data, we aim to establish a reproducible framework using multimodal parameterization methods.

**Methods:** Herein we developed physiologically based kinetic (PBK) models, following oral exposure. Their parametrization was primarily based on structural, physiological and experimental values, as well as quantitative structure-activity relationship (QSAR) predictions. Outputs were evaluated against available biomonitoring data for BPA and BPS. Critical parameters were identified by sensitivity analysis and iteratively re-sampled in Monte Carlo (MC) simulations to quantify uncertainties.

**Results:** Among human models parametrized for males and females of different ages, we predicted that bisphenols reached the highest concentrations in 5-year-old males. Environmentally relevant exposure levels resulted in maximum concentrations in the blood and testes for BPS, and in the thyroid for BPM. After 96 hours, steady-state concentrations were not yet reached in the breasts for BPA, BPAF, BPB, BPE, BPF and BPM.

**Conclusions:** The data from this study suggest significant variability in internal concentrations for identical exposures to different bisphenols analogs that further depend on age, sex and organ. This diversity in toxicokinetic behavior should be considered for health risk assessment of these substitutes.

## Introduction

The rigid molecular structure of bisphenol compounds confers essential material properties for plastic production, such as chemical resistance and thermal stability.^1^ In numerous countries, including Canada,^2^ China, Japan, Korea,^3^ the United States (US)^4^ and the European Union,^5,6^ bisphenol A (BPA), once an industry staple, now faces restrictions or bans, leading to an increasing use of its structural analogs as substitutes. BPAF, BPB, BPE, BPF, BPM and BPS were detected in beverage packaging material, some of them in BPA-free labeled products.^7^ Bisphenols substitutes can migrate from packaging materials to their content,^8^ which raises significant concerns, as their structural similarity to BPA suggests they might also exert endocrine disrupting (ED) effects by binding to biological receptors, such as estrogen, androgen, and thyroid hormone receptors.^3,9^ These interactions can lead to adverse outcomes, notably, reproductive disorders, thyroid and gonadal dysfunction/cancer, breast cancer, obesity and diabetes.^10^ Indeed, BPB and BPS have been identified as a substance of very high concern under the European REACH Regulation due to their ED properties and reproductive toxicity for BPS.^11,12^ The opinion of ECHA’s committee for risk assessment on BPAF as toxic to reproduction (Repr. 1B) is in the European Commission for decision-making^13^ and this compound was also found to meet the ED criteria for human health.^14^ To evaluate the safety of BPAF, BPB, BPE, BPF, BPM and BPS, Next Generation Risk Assessment (NGRA) principles recommend an exposure-led approach based on internal exposure levels.^15^ This involves converting external exposure data into concentration-time profiles in blood and organs, underscoring the necessity of prior knowledge of absorption, distribution, metabolism, and excretion (ADME) processes to determine relevant points of departure for hazard predictions. Since human internal exposures of these six bisphenol analogues are not yet characterized, their risk assessment remains largely hindered.

Alterations in human exposure patterns to bisphenols were found to coincide with shifts in industrial usage. In three studies of urine concentrations of bisphenols in the US and Japan between 1993 and 2019, there was a decline in BPA levels alongside a rise in BPE, BPF and BPS levels.^16–18^ In Czech healthy normospermic men, BPF plasma levels were three times higher than BPA levels.^19^ BPS was detected most frequently in breastmilk from breastfeeding women in south China^20^ and was found in over 80% of semen samples of Czech men.^21^ In China, BPAF was detected in 28% of urine samples om pregnant women, and was suspected to influence the homeostasis of thyroid-related hormones in cord blood.^22^ BPB was identified as the fourth most prevalent bisphenol in urine samples of Flemish adolescents.^23^ In Swiss infants and toddlers aged between 6 and 36 months, BPM reached the highest urinary concentration (8.56 µg/g) among BPA and 14 emerging BP analogues.^24^ It was also frequently detected in the saliva of both obese and non-obese Spanish children between 6 and 12 years old.^25^ These data confirm that the widespread use of bisphenol analogs results in important human exposure, particularly among vulnerable populations, yet fail to provide essential toxicokinetic insights.

The kinetic behavior of BPA following oral administration has been well characterized.^26,27^ After a rapid and efficient absorption in the gastrointestinal tract, its phase II conjugation in the gut wall and the liver leads to the extensive production of its main metabolite, BPA-glucuronide.^26^ It is assumed that other bisphenols also undergo a similar metabolization pattern forming biologically inactive conjugated metabolites.^28^ BPA excretion occurs from the blood via the kidneys and is rapidly eliminated via urine.^26^ To predict chemical concentrations in key organs, following various exposure scenarios, further mechanistic insights were provided by physiologically based kinetic (PBK) models. An eight-compartment PBK model capable of predicting human internal BPA exposures was proposed by Yang et al. (2015). This model is largely calibrated from serum time-concentration profiles measured in a biomonitoring study.^30^ Since even slight differences in xenobiotic structures can induce significant variations in ADME processes, BPA analogs cannot be assumed to share identical kinetics, making it essential to evaluate them all. However, while the toxicokinetics of BPS following oral exposure have been studied,^31^ to the best of our knowledge, BPAF, BPB, BPE, BPF, and BPM have not been similarly assessed in humans. The lack of data limits the application of the BPA PBK model parametrization method^29^ to other bisphenols, and thus, the prediction of their internal concentrations in key organs.

Karrer et al. (2018) addressed the challenge of extending the existing BPA PBK model to BPS, BPAF, and BPF. Their approach relied on the molecular weight (MW) and structural similarities: BPA and BPF were grouped together, as were BPS and BPAF, to allow the re-use of biomonitoring data from well-characterized bisphenols for calibrating models of data-poor analogues. Although this model solves the issue of extending predictions to less-characterized bisphenol substitutes, its application remains limited to bisphenols with minimal physico-chemical differences from BPA and BPS.

To address the lack of human toxicokinetic data for a wider diversity of bisphenols, we constructed PBK models for BPA and six of its structural analogs using the OECD integrative parametrization and harmonized methodology (Figure S1).^33^ We hypothesized that these models could predict their internal exposures following oral administration, achieving a level of accuracy comparable to that of previous models developed with calibration to human data.^31,32^ A common concept was established for four standard human models, varying by age and sex. The models for each bisphenol analog were individually parametrized. Physico-chemical properties were predicted from chemical structures. Intestinal uptake and blood-to-tissue partition coefficients were obtained using quantitative structure-activity relationship (QSAR) methods and quantitative *in vitro* to *in vivo* extrapolation (QIVIVE). Hepatic and intestinal glucuronidation kinetics were measured in S9 fractions. Enterohepatic circulation (EHC) was parameterized based on rat PBK models predictions. Following parametrization, the model performance was evaluated against human *in vivo* concentrations of BPA and BPS and compared to the outcomes of previous modeling efforts. Concentrations in twelve compartments including blood, thyroid, testes and breasts were predicted after environmentally relevant oral exposure of BPA, BPAF, BPB, BPE, BPF, BPM and BPS. Sensitivity analyses were conducted to identify critical compartments and parameters, which were then resampled in 10,000 Monte Carlo (MC) simulations to account for interindividual variation and uncertainty.

## Methods

### Scope and purpose of exposure scenarios

Diet is the main route of human exposure to bisphenols, therefore models were designed herein for oral administration.^34^ Several exposure scenarios were defined for parameter gathering, model evaluation and internal exposure assessment (Table 1). In scenario 1 and 2, a rat PBK model was used to parameterize EHC rates for application in human PBK models. Scenarios 3 and 4 replicated the administration of BPA from Teeguarden et al. (2015) and BPS from Oh et al. (2018) for model evaluation. To illustrate potential BPA replacement by its analogues: BPAF, BPB, BPE, BPF, BPM and BPS, scenarios 5 to 12 assumed consistent exposure rates amongst all these bisphenols, based on the 2015 EFSA assessment of BPA external exposure.^35^ Four physiological human models were used: adult male or “man” (body weight (BW): 73 kg, height (H): 176 cm, age: 30 years), adult female or “woman” (BW: 60 kg, H: 163 cm, age: 30 years), child male or “child” (BW: 19 kg, H: 109 cm, age: 5 years), and toddler male or “toddler” (BW: 10 kg, H: 76 cm, age: 1 year). These models were selected based on available physiological data for key organs (thyroid, breasts, and testes), related to ED and detected bisphenol analog exposure level. The repeated dose scheme from Karrer et al. (2018) was reproduced (scenarios 9 to 12) to model chronic exposure via three meals a day.

**Table 1.** Exposure scenarios, scope and purpose of modeling.

| scenario | purpose | model | BP | dosing | key organ(s) |
| --- | --- | --- | --- | --- | --- |
| 1 | EHC determination | rat | BPAF | single dose<sup>a</sup>: 340 mg/kg bw | blood |
| 2 | EHC determination | rat | BPF | single dose<sup>a</sup>: 200 mg/kg bw | blood |
| 3 | model evaluation | man<sup>b</sup> | BPA | single dose<sup>a</sup>: 30 µg/kg bw | blood |
| 4 | model evaluation | adult<sup>c</sup> | BPS | single dose<sup>a</sup>: 8.1 µg/kg bw | blood |
| 5 | internal exposure assessment | man | <sup>d</sup> | single dose<sup>a</sup>: 336 ng/kg bw | blood, thyroid, testes |
| 6 | internal exposure assessment | woman | <sup>d</sup> | single dose<sup>a</sup>: 389 ng/kg bw | blood, thyroid, breasts |
| 7 | internal exposure assessment | child | <sup>d</sup> | single dose<sup>a</sup>: 818 ng/kg bw | blood, thyroid, testes |
| 8 | internal exposure assessment | toddler | <sup>d</sup> | single dose<sup>a</sup>: 869 ng/kg bw | blood, thyroid, testes |
| 9 | internal exposure assessment | man | <sup>d</sup> | repeated dose<sup>e</sup>: 112 ng/kg bw | blood, thyroid, testes |
| 10 | internal exposure assessment | woman | <sup>d</sup> | repeated dose<sup>e</sup>: 130 ng/kg bw | blood, thyroid, breasts |
| 11 | internal exposure assessment | child | <sup>d</sup> | repeated dose<sup>e</sup>: 273 ng/kg bw | blood, thyroid, testes |
| 12 | internal exposure assessment | toddler | <sup>d</sup> | repeated dose<sup>e</sup>: 290 ng/kg bw | blood, thyroid, testes |

<sup>a</sup> Single oral dosing at t=0 h. Total duration of the simulation: 48 h.
<sup>b</sup> BW and H adjusted to the population (10 men) of the biomonitoring study by Teeguarden et al. (2015).
<sup>c</sup> BW and H adjusted of the population (4 men and 3 women) of the biomonitoring study by Oh et al. (2018).
<sup>d</sup> BPA, BPAF, BPB, BPE, BPF, BPM and BPS.
<sup>e</sup> Repeated oral dosing at t=0, 6, 12, 24, 30, 36,48, 56, 60,72, 78 and 84 h. Total duration of the simulation: 96 h.
Note: BP, bisphenol; BPA, bisphenol A; BPAF, bisphenol AF; BPB, bisphenol B; BPE, bisphenol E; BPF, bisphenol F; BPM, bisphenol M; BPS, bisphenol S; EHC, enterohepatic circulation; BW, body weight; H, height.

### PBK model structure

A common model concept was employed for the seven bisphenols (Figure 1). Oral administration was modeled through the stomach compartment, followed by gastric emptying into the gut lumen and absorption into gut tissue. Upon reaching the liver, each bisphenol was modeled to undergo glucuronidation, and changes in concentration of the respective metabolite over time were further simulated in metabolite specific compartments. To represent EHC, we modeled a portion of hepatic glucuronidated bisphenol to be cycled back through the bile into the gut lumen, where it was expected to revert to its parent form. Non-glucuronidated hepatic bisphenols were simulated to reach systemic circulation, including distribution to blood-perfused organs. When not individually compartmentalized, these organs were categorized as either slowly perfused tissues (e.g., muscles, bone) or rapidly perfused tissues (e.g., heart, lung, kidneys) tissues and combined. Compartments related to the thyroid and testes/breasts (depending on sex) were included for the human, but not the rat PBK models. The model was developed using Berkeley Madonna 10 and Python. To address the challenge of transferability across different modeling platforms, both scripts are provided (supporting information and on GitLab : https://gitlab.ethz.ch/eth_toxlab/pbk_model_bisphenol_analogs).

**Figure 1.**
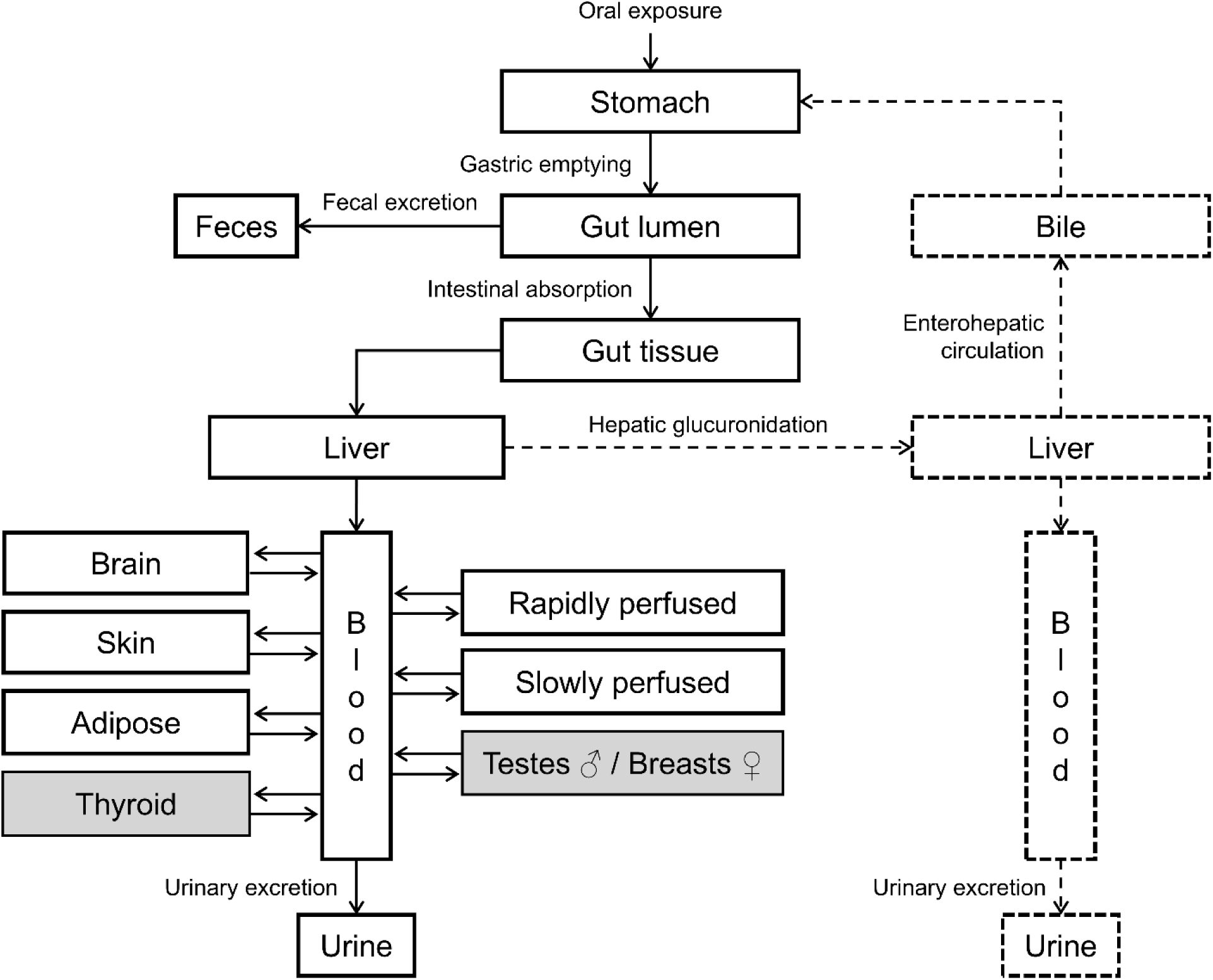
Schematic PBK model concept for BPA analogs (flows and compartments with continuous lines) and their glucuronidated metabolites (flows and compartments with dashed lines). Compartments with grey filling are only present in human models. Note: ♂, male; ♀, female; BPA, bisphenol A; PBK; physiologically based kinetic.

### Model parameterization

Molecular parameters were adjusted for each bisphenol, including both original and glucuronidated forms (Figure 2). Parameters related to physico-chemical characteristics of the seven modeled bisphenols and their glucuronidated compounds were predicted from their structure with ChemDraw 20.0 software (see Table 2 and Table S1). These data were also used to predict intestinal uptake and partition coefficients.

**Figure 2.**
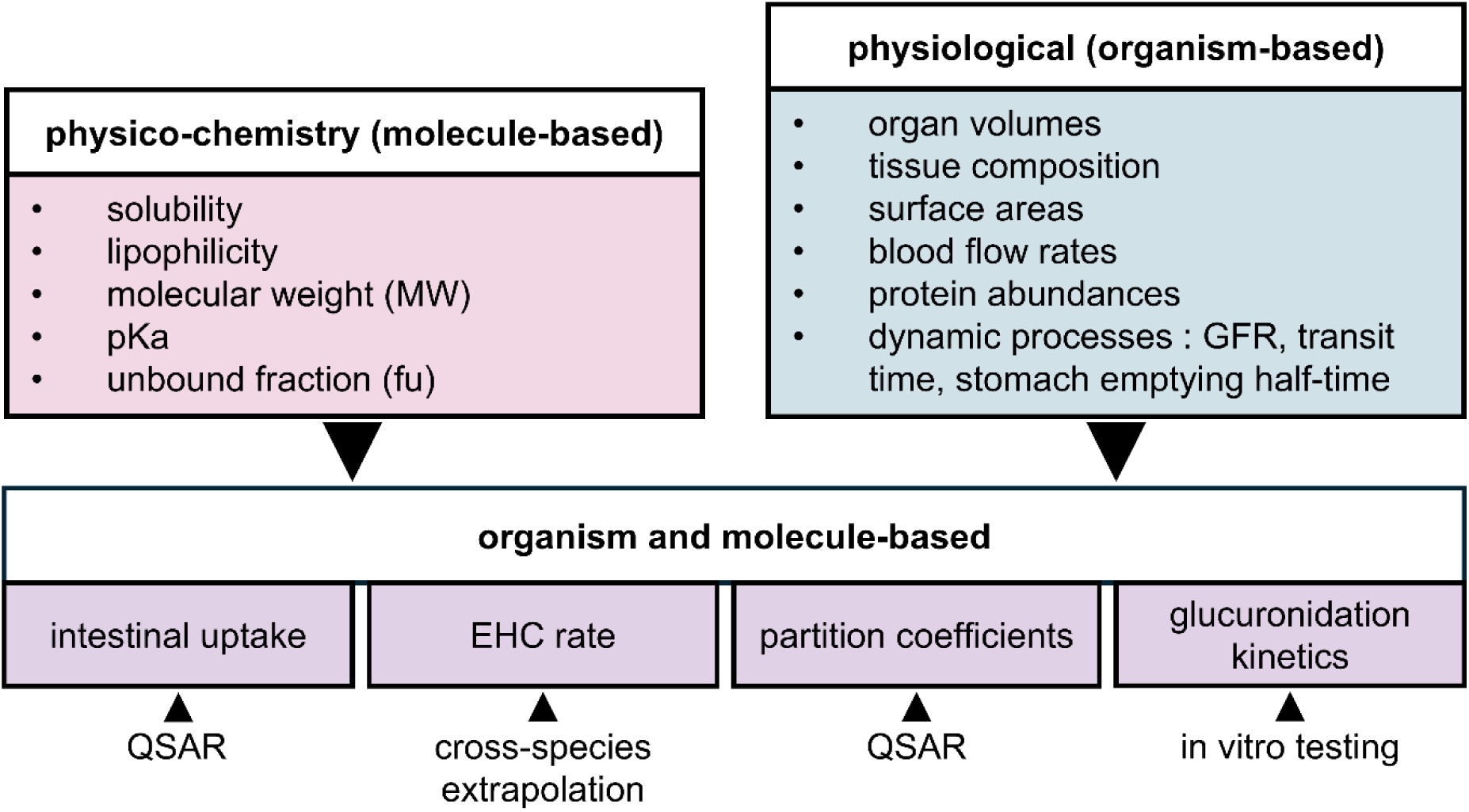
Overview of parameters used in the present model and how they were acquired. Note: GFR, glomerular filtration rate; QSAR, quantitative structural-activity relationship; EHC, enterohepatic circulation.

**Table 2.**
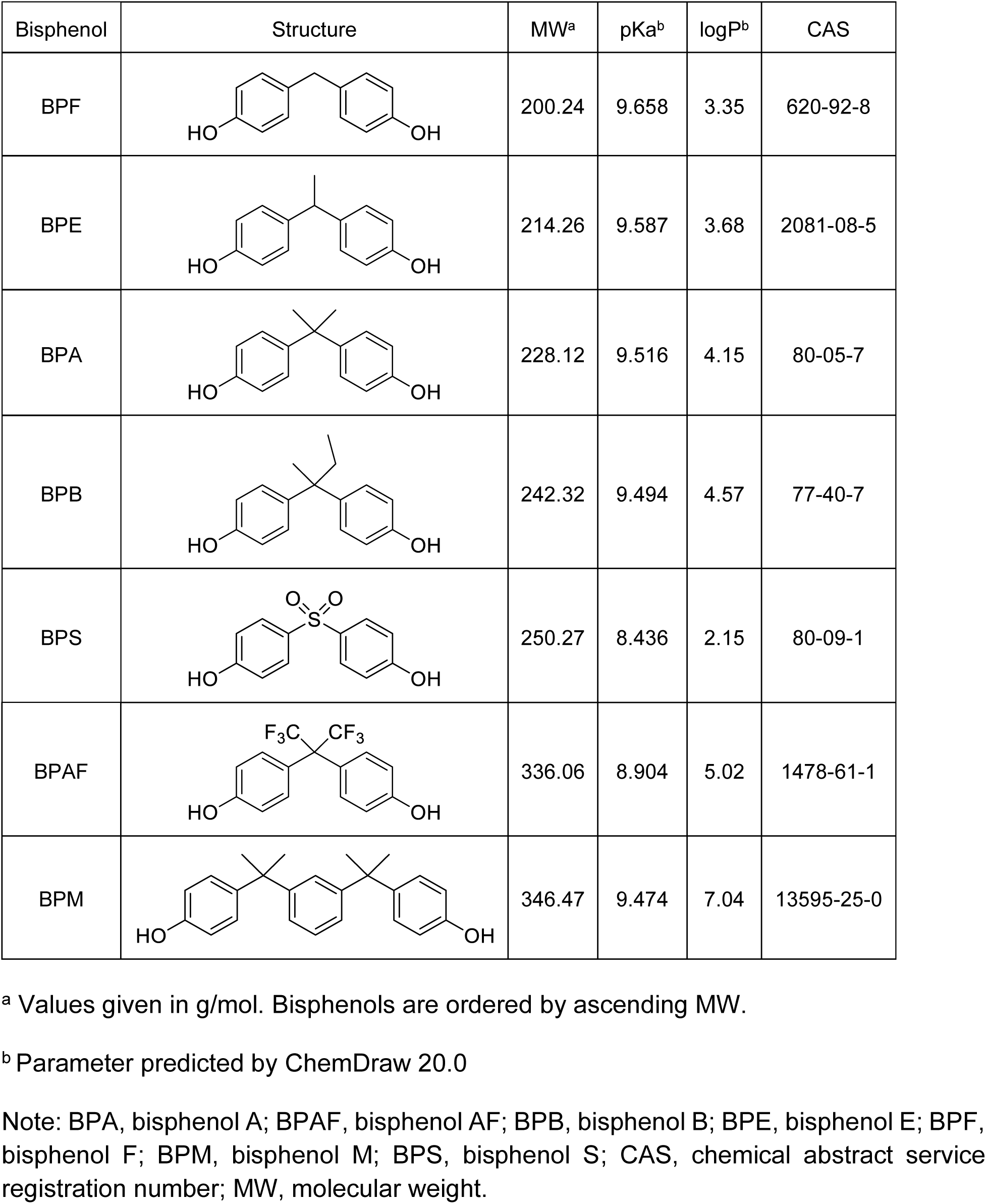
Physico-chemical properties of bisphenol analogs included in the present study.

| Bisphenol | Structure | MW<sup>a</sup> | pKa<sup>b</sup> | logP<sup>b</sup> | CAS |
| --- | --- | --- | --- | --- | --- |
| BPF |  | 200.24 | 9.658 | 3.35 | 620-92-8 |
| BPE |  | 214.26 | 9.587 | 3.68 | 2081-08-5 |
| BPA |  | 228.12 | 9.516 | 4.15 | 80-05-7 |
| BPB |  | 242.32 | 9.494 | 4.57 | 77-40-7 |
| BPS |  | 250.27 | 8.436 | 2.15 | 80-09-1 |
| BPAF |  | 336.06 | 8.904 | 5.02 | 1478-61-1 |
| BPM |  | 346.47 | 9.474 | 7.04 | 13595-25-0 |

<sup>a</sup> Values given in g/mol. Bisphenols are ordered by ascending MW.
<sup>b</sup> Parameter predicted by ChemDraw 20.0
Note: BPA, bisphenol A; BPAF, bisphenol AF; BPB, bisphenol B; BPE, bisphenol E; BPF, bisphenol F; BPM, bisphenol M; BPS, bisphenol S; CAS, chemical abstract service registration number; MW, molecular weight.

Organism-based parameters were tailored to account for species, sex, and age when possible (Figure 2, Table S2).

Human physiological parameters were predominantly sourced from the International Commission on Radiological Protection (ICRP) report 89.^36^ Fractional organ volumes were computed with the formula (1), except for breasts volume, which was the ratio between breasts and whole-body weight. The volumes of combined compartments were the sum of their constituent organ volumes. Concerning gut tissue fractional volume, the total mass of gastrointestinal tract walls (excluding esophagus and stomach) was divided by body weight, while the fractional volume of gut lumen was estimated by dividing the total mass of gastrointestinal tract content (excluding the stomach) by body weight. As described by Helander and Fändriks (2014), total intestinal surface area (ISA) was modeled by approximating the small intestine (SI) and large intestinal (LI) as cylinders enlarged with folds, villi and microvilli.^37,38^ Length values for SI and LI were adjusted based on age and sex, while diameter values from Helander and Fändriks (2014) were the same for all human models. Blood flow rates (as % of cardiac output) were set from adult values. Lung blood flow rate issued from Willmann et al. (2007). The gut tissue blood flow rate was the sum of SI and LI blood flow rate. The blood flow rates of combined compartments were the sum of their constituent organs blood flow rates. Gastric emptying half-time was measured in Oberle et al. (1990) for an administered volume of 200 mL, to approach the conditions of the biomonitoring study.^30^ The intestinal transit time was defined as the sum of the transit times of luminal contents through SI, right colon, left colon, and sigmoid colon. Human glomerular filtration rate (GFR) was sourced from Levey et al. (2014).

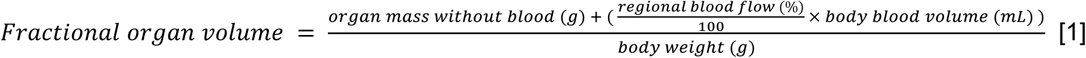

Rat physiological parameters were primarily sourced from Brown et al. (1997), to represent an eleven-week-old Sprague Dawley rat weighing 250g, with no specification of sex. Stomach and gut tissue (intestine) volume were measured by Oatley and Toates (1969) in hooded rats. Gut lumen volume was the sum of fluid volumes in the duodenum, upper jejunum, lower jejunum and ileum for Wistar rats following oral administration of 1 mL.^44^ Intestinal surface area value was sourced from Meshkinpour et al. (1981). Gut tissue blood flow rate (as % of cardiac output) was sourced by Davies and Morris (1993). The blood flow rates of combined compartments were the sum of their constituent organs blood flow rates. The gastric emptying half-time for a volume of 1 mL was extracted from Purdon and Bass (1973) to approach the volume delivered in animal study (1.25 mL, Waidyanatha et al. (2019)). Intestinal transit time was the sum of the values reported for chyme transit times through SI and LI.^49^ The rat GFR value was sourced from Davies and Morris (1993).

Parameters based on both organism and molecule were partition coefficients, intestinal uptake, EHC rate and glucuronidation kinetics (Figure 2). The first three were obtained from *in silico* approaches.

Fo most compartments, the same partition coefficients and plasma unbound fractions were used for different ages, sex and species (Supplementary Table S3). Both were calculated with the QIVIVE toolbox, using the methods of Rodgers and Rowland (2006) and of Lobell and Sivarajah (2003).^52^ Partition coefficients for combined compartments were the weighted sums of partition coefficients for the constituent organs. Thyroid and testes partition coefficients were calculated using data from Pilari et al. (2017). Based on Ulaszewska et al. (2012), breasts partition coefficients were calculated as the product of adipose partition coefficients and breasts fat volume, which was set to 24.5% by averaging data from Vandeweyer and Hertens (2002).

To parametrize the intestinal uptake (k_a_) of all bisphenols, the Caco-2 apparent permeability coefficient (P_app_) was converted into intestinal effective permeability (P_eff_), and then scaled to the whole body with ISA value. P_app_ values were only available for BPA, BPF, and BPS.^56^ Additionally, two QSARs were used to predict P_app_ values for the seven bisphenols. The first, “QSAR K” from Kamiya et al. (2020), was trained on a data set including the P_app_ for BPA, BPF and BPS. LogD (lipophilicity) values, required for these predictions, were computed using the Chemaxon predictor,^57^ considering pH values of 6.0 (apical) and 7.4 (basal). The second, “QSAR L”s an approach developed by Lanevskij and Didziapetris (2019), which computes permeability from the inverse of overall resistance to diffusion. It was trained on a dataset of 1366 P_app_ values that did not include any bisphenol. For these predictions, standard values were set for pH (7.4), stirring rate (0 rpm), and average pore radius (5 Å), while McGowan characteristic volumes were calculated according to Zhao et al. (2003). Then, these two (BPAF, BPB, BPE, BPM) or three (BPA, BPF, BPS) P_app_ values were averaged and the scaling algorithm of Sun et al. (2002) was used (with the regression for all drugs at pH 7) to obtain the P_eff_ of each bisphenol analog. Then, for each bisphenol and physiological model, k_a_ was the product of P_eff_ and ISA.

EHC of all glucuronides was modeled by a rate between 0 and 1, representing the portion of metabolites that undergo biliary excretion. This value was optimized by fitting model predictions of blood concentration at late time points, as recently described by Aichinger et al. (2023). Available plasma concentrations of BPAF^48^ or BPF^62^ in rats following a single oral dose were used for this purpose. Predicted-to-observed ratios of maximum concentrations (C_max_) in blood were calculated. The EHC rate, initially set to zero, was incrementally adjusted until predictions of blood concentrations at 24 hours (BPF) and 48 hours (BPAF) fit the observed values, whilst conserving the blood C_max_ ratio. Given the importance of MW for EHC,^63^ with BPAF being the lightest and BPF the second heaviest bisphenol in this study, EHC values for other bisphenols were calibrated by MW. For simplicity, the EHC of BPM was set equal to BPAF. Considering the stability of the relative expression of glucuronide biliary excretion transporter MRP2 (Multidrug resistance protein 2) for all ages concerned by the present study,^64^ the same values were used all PBK models for humans of the same sex.

### Measurement and parametrization of glucuronidation kinetics

The glucuronidation kinetics of BPA, BPB, BPE, BPF, BPAF, and BPM were determined as follows (Table S4): BPA, BPB, BPE and BPM were incubated with Corning® UltraPool™ Human Liver S9 in a buffer containing UDPGA. The S9 reagent (20 mg protein/mL) was aliquoted and stored at −80° C until use. BPAF and BPF were incubated similarly with rat liver S9 (Sprague-Dawley, male, 0.1 mg/mL S9 protein). For each assay, an S9 aliquot was thawed on ice, and a reaction master mix was prepared with 0.1 mg S9 protein/mL in either 0.1 M (for BPA, BPB, BPAF, and BPF) or 0.25 mg/mL (for BPE and BPM) Tris-HCl, along with 10 mM UDPGA in 50 mM Tris-HCl (pH 7.4), 25 µg/mL alamethicin, and 10 mM magnesium chloride. Negative controls using 0.1 M Tris-HCl without UDPGA were included. Each reaction solution (100 µL) was pre-incubated for 5 min at 37° C with 350 rpm shaking. To this solution was added bisphenol solutions (0 to 20 mM) in DMSO in a ratio of 1:100 to achieve final bisphenol concentrations of 0 to 200 µM and 1% (v/v) DMSO. The shortest incubation times for maximal reaction speeds were identified: 10 min for BPE, 15 min for BPB, 20 min for BPA and BPM, 60 min for BPF, and 120 min for BPAF. The glucuronidation reaction was stopped by adding cold methanol (160 μL) and samples were kept at −20 °C for 1 h to precipitate proteins. The resulting mixture was centrifuged (10 min, 4 °C, 15,000 rcf). Supernatants were analyzed using High-performance liquid chromatography (HPLC) with diode array detector (DAD). Similar experiments were conducted for BPA, BPB, BPE, and BPM with human small intestinal S9 mix (0.025 to 0.25 mg/mL). Bisphenol concentrations ranged from 5 to 120 µM, and incubation times from 15 to 120 min.

The quantification of the bisphenols and their glucuronides was carried out on an Agilent 1200 series HPLC system with a Waters XBridge BEH130 C18 column (4.6 × 150 mm, 3.5 μm particle size) for separation and a DAD set to a detection wavelength of λ = 280 nm. A gradient elution was used with water containing 0.1% formic acid as eluent A and acetonitrile with 0.1% formic acid as eluent B. The initial condition of 15% eluent B was increased to 65% eluent B over 20 min, and then to 90% eluent B in the next 5 min. The retention times for the various bisphenols and their glucuronides are provided in Supplementary Table S5. Calibration curves were set from concentrations ranging from 1 to 100 µM.

Data analysis of glucuronidation kinetics included exclusion of the outliers using the Nalimov test.^65^ Glucuronidation rates were graphically determined based on compound loss over time using GraphPad Prism 10.2.3 (2024). Michaelis-Menten and substrate inhibition curve fitting algorithms were applied, and the best fit, indicated by maximal R², was selected.

Experimental glucuronidation rates were scaled using specific factors based on the *in vitro* context from which they were obtained. Thus, data from human hepatic S9 fractions was scaled with 107.3 mg S9 protein per gram liver,^66^ data from rat hepatic S9 fractions with 143 mg/g liver,^67^ and data from human hepatic microsomes with 32 mg protein/g liver.^32,68^ To account for age-related variation in expression levels of the UDP glucuronosyltransferase UGT2B15 protein, an ontogeny-scaling method was used to establish scaling factors for glucuronidation (SF_g_) kinetics (Equation 2).^69^ Protein abundances were derived from Bhatt et al. (2019), with both child and toddler models classified under early childhood.

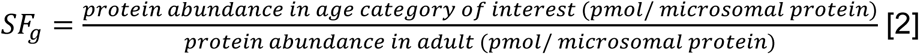

### Model evaluation

The performance of rat and human PBK models was evaluated by comparing predicted blood C_max_, t_max_, and AUC values to their *in vivo* measured values, for BPAF,^48^ BPF,^62^ BPA^30^ and BPS.^31^ Models were refined until predicted blood concentration where similar to corresponding experimentally observed values.^33^ Toxicokinetic parameters predicted for BPA, BPA glucuronide, BPS and BPS glucuronide were compared to previous modeling results.^31,32^

### Sensitivity analysis

The impact of parameter deviation on model predictions was assessed by a sensitivity analysis based on the method by Evans and Andersen (2000). For this, in all human models and all bisphenols, parameters were individually increased by 5% and the associated impact on blood, thyroid and breast or gonad concentrations was computed. Normalized sensitivity coefficients (SC) were determined by using Equation 3. P refers to the value of the unchanged parameter of interest and P’ to its elevated value. C_max_ refers to the maximal concentration of bisphenol predicted in the compartment with unchanged values of parameters and C_max_’ refers to the same with increased value of the parameter. A parameter with an absolute SC value > 0.1 was considered sensitive.

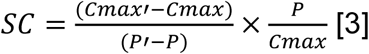

### Uncertainty analysis

To address parameter uncertainty in concentrations that were predicted with human PBK models, 10,000 MC simulations were performed. Each sensitive parameter was re-sampled according to probability distributions from the literature (Table S6)^39,40,70,72–76^. Coefficients of variation were calculated with the ratio of standard deviation (σ) and mean (μ) for standard measurements, or with equation 4 for ratios (e.g., SFg). To ensure physiological plausibility, EHC rates were constrained between 0 and 1, and the sums of all organs fractional blood flood rates and all organs fractional volumes were conserved during resampling. MC simulation outcomes were analyzed by comparing first quartile, median, and third quartile values of concentrations for each time point.

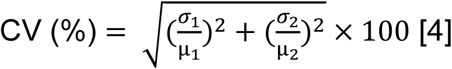

## Results

### Parametrization of intestinal uptake

To develop PBK models for predicting internal concentrations of BPA and six of its structural analogs following oral administration, it was necessary to accurately parameterize their absorption into the gut tissue. We compared P_app_ values of the bisphenols analogs, either measured in Caco-2 cells monolayers (BPA, BPF and BPS) and/or predicted by two QSARs (all analogs) in Figure 3.^56,58^ For BPA, BPAF, BPB, BPE, and BPF, all various techniques yielded similar results, ranking the permeability in the same order, with values differing by a maximum of 15×10^−6^ cm/s. This consensus was not observed, however, for BPM and BPS. Among BPM predictions, QSAR L (considering molecule volume) predicted lower permeability than was predicted with QSAR K. For BPS, the two QSAR predictions of P_app_ values were close to each other, but they were much lower than the corresponding data from measuring transport *in vitro*. Additionally, BPS and BPAF, which contain substituents with higher electronegativity (sulfur and fluor), were the only bisphenols where QSAR K predictions were lower than QSAR L predictions. For the three bisphenols where comparison was possible (BPA, BPF and BPS), both QSARs predicted P_app_ values with a similar level of proximity to the *in vitro* measurements, indicating comparable performances. These results demonstrate that the parameterization of intestinal uptake for each bisphenol analog is possible without the need for read-across and provide insights into potential biases of *in silico* methods.

**Figure 3.**
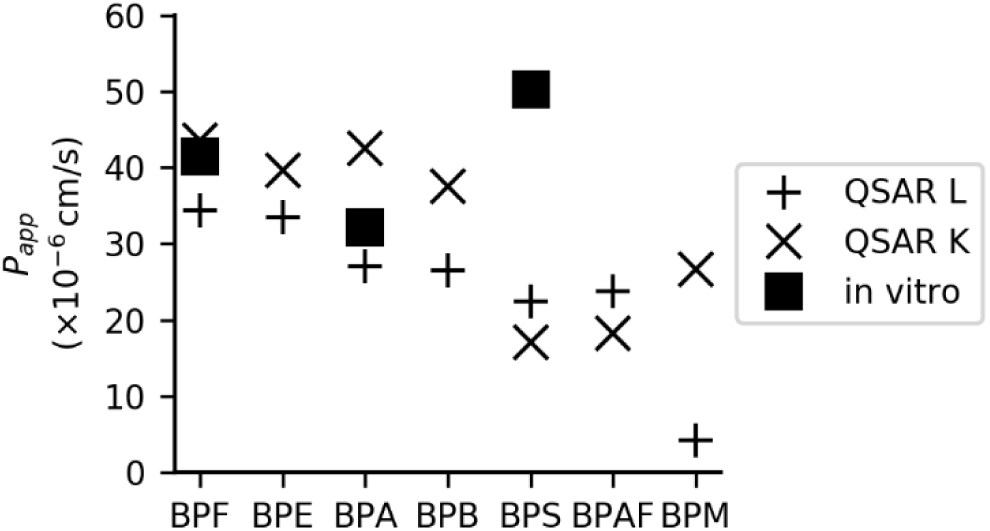
Estimated P_app_ across the Caco-2 monolayer determined by various techniques. Squares are results from *in vitro* experiments with BPF, BPA and BPS,^56^ while crosses refer to predictions for all seven bisphenols by two different QSARs (K: Kamiya et al. (2020) and L: Lanevskij and Didziapetris (2019)). Bisphenols are ordered by ascending MW. Note: BPA, bisphenol A; BPAF, bisphenol AF; BPB, bisphenol B; BPE, bisphenol E; BPF, bisphenol F; BPM, bisphenol M; BPS, bisphenol S; P_app_, apparent permeability coefficient, QSAR, quantitative structural activity relationship.

### Parametrization of hepatic glucuronidation kinetics

To parametrize bisphenol metabolism processes occuring in the liver compartment, we measured hepatic glucuronidation kinetics of BPA, BPB, BPE, and BPM in human liver S9 fractions) and included previous measurements in hepatic microsomes for BPS, BPF, and BPAF (Figure 4, Supplementary Figure S2, Supplementary Table S7. Among the analogs, BPB was hepatically glucuronidated the most efficiently, while BPS was the least. Negligible intestinal metabolism rates were measured for BPA, BPB, BPE, and BPM. Indeed, the highest observed glucuronidation speed was 0.2 nmol per min per mg S9 protein, corresponding to the incubation of 50 µM BPE with 0.125 mg/mL intestinal S9 mix for 90 min. Hepatic glucuronidation of BPAF and BPF were measured in rat liver S9 fractions for the parameterization of the rat model (Supplementary Figure S2).

**Figure 4.**
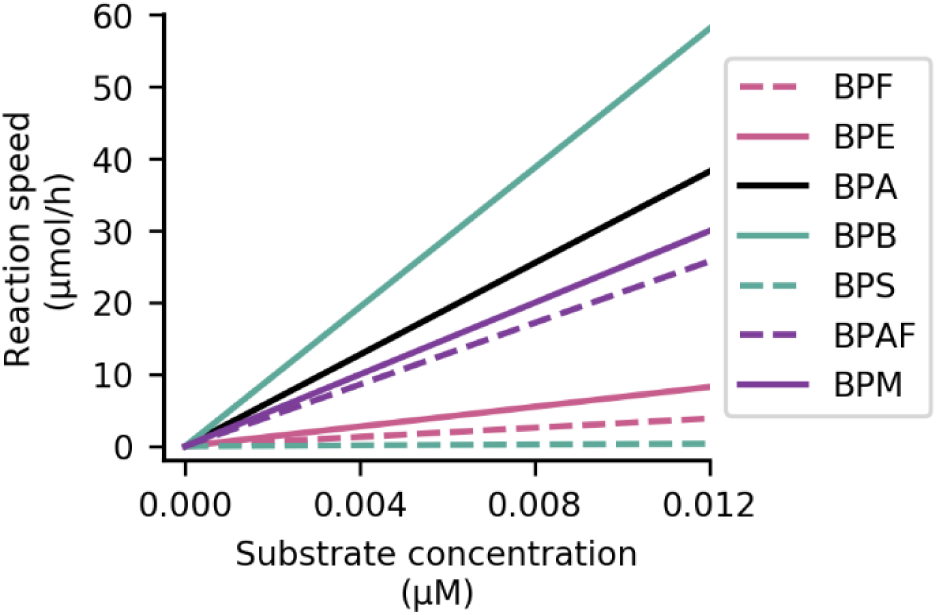
Predicted rates of glucuronidation of bisphenols in human adult liver at physiologically relevant concentrations. Kinetics of glucuronidation catalyzed by S9 fractions (BPA, BPB, BPE, and BPM, continuous lines) or microsome fractions (BPS, BPF and BPAF, Karrer et al. (2018), dashed lines). Michaelis-Menten kinetics were used to describe the glucuronidation of BPA, BPF and BPS, while substrate-inhibition kinetics were used for BPB, BPE, BPM and BPAF. Bisphenols are ordered by ascending MW. Note: BPA, bisphenol A; BPAF, bisphenol AF; BPB, bisphenol B; BPE, bisphenol E; BPF, bisphenol F; BPM, bisphenol M; BPS, bisphenol S.

### Parametrization of biliary excretion

Since EHC is a crucial process for the residence time and accumulation of xenobiotics, its inclusion in the PBK models was considered essential for accurate predictions of bisphenol analog concentrations over time. Thus, EHC rates were introduced by incrementally adjusting this parameter in rat models until the predicted blood concentrations matched *in vivo* measurements at later time points, and these values were then directly used in human models (Table 3). Thereby, a similar high EHC rate was predicted for each bisphenol, with more than 73% of glucuronidated bisphenols leaving the liver though biliary excretion. Regarding sex differences, EHC values were slightly higher in females due to higher BPAF concentrations measured in female rats 24 h post-administration.^48^ Additionally, the half-life of BPF was longer in female rats compared to males.^62^

**Table 3.** EHC rate values. Bisphenols are ordered by ascending MW.

| Sex | BPF | BPE | BPA | BPB | BPS | BPAF | BPM |
| --- | --- | --- | --- | --- | --- | --- | --- |
| Male | 0.73 | 0.74 | 0.74 | 0.75 | 0.76 | 0.80 | 0.80 |
| Female | 0.73 | 0.75 | 0.76 | 0.78 | 0.79 | 0.88 | 0.88 |

Note: BPA, bisphenol A; BPAF, bisphenol AF; BPB, bisphenol B; BPE, bisphenol E; BPF, bisphenol F; BPM, bisphenol M; BPS, bisphenol S.

### PBK models evaluation with BPA and BPS biomonitoring

The present model was evaluated with measures from Teeguarden et al. (2015), and predicted blood C_max_ values for BPA and its glucuronide were 53% and 72% of the observed values, respectively (Figure 5). In the present predictions, t_max_ for BPA was lower than the observed value, while t_max_ for BPA glucuronide was similar. Predicted AUC for BPA closely matched the *in vivo* value, while predicted AUC for BPA glucuronide was about 2 times higher than observed. Five of six predicted values were within the measurement ranges, and the variability predicted by MC simulations overlapped with all measured ranges. These results indicate that the present model generates reliable predictions of BPA concentrations. Furthermore, five of the six toxicokinetic parameters values predicted by the present model were closer to the measures than values predicted by previous models, which suggests superior performances of the present model for BPA concentrations predictions.^31,32^

**Figure 5.**
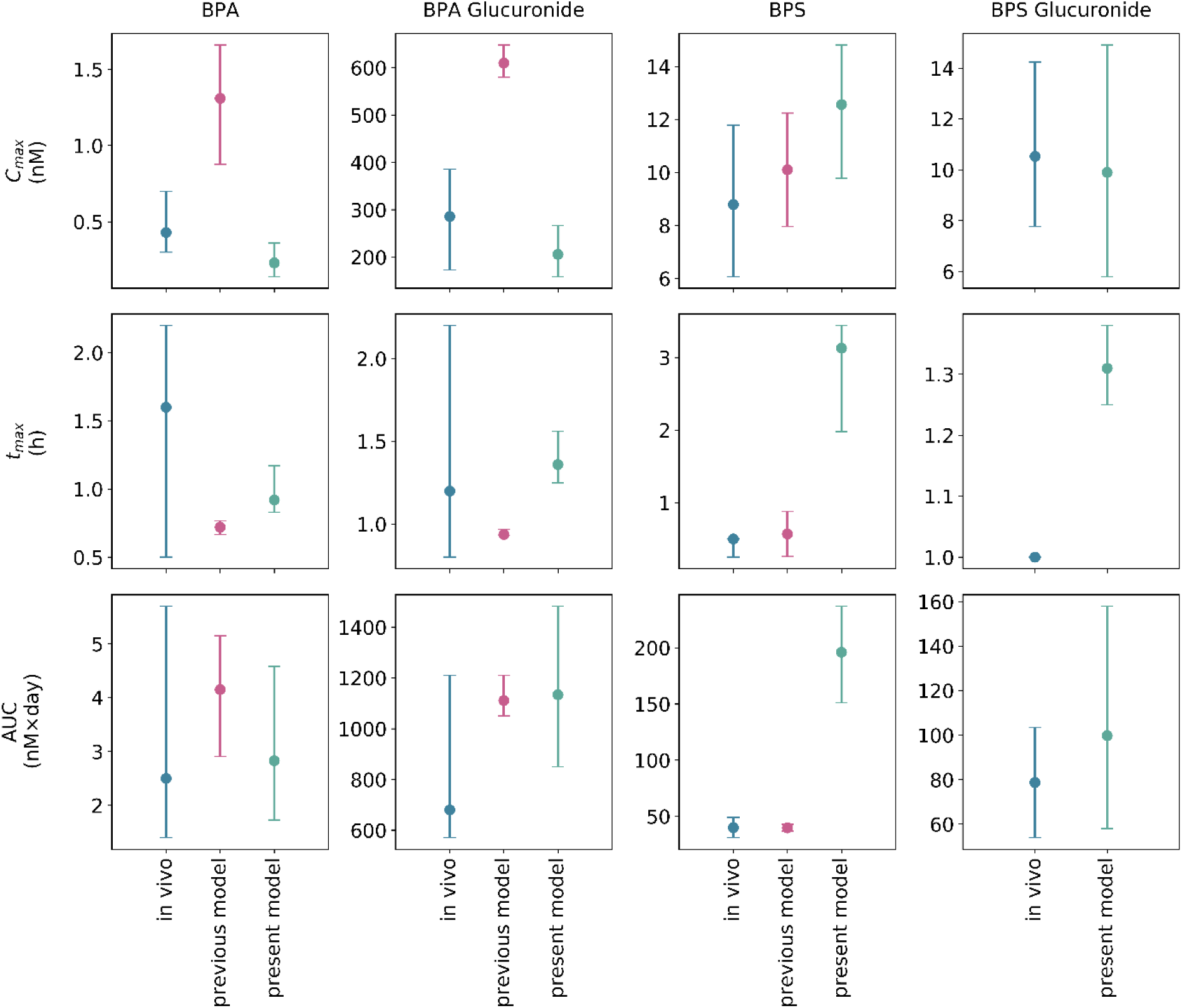
Comparison of measures and predictions of toxicokinetic parameters for oral ingestion of BPA (left) or BPS (right). Measured toxicokinetic parameters values (“in vivo”) are in blue, toxicokinetic parameters predicted by previous modeling (“previous model)” in grey and toxicokinetic parameters predicted by present modeling in pink. For measurements and results from previous models, errors bars correspond to reported values and prediction range.^31,32^ Intervals of variability of the present model predictions were set as first and third quartiles of the MC simulations. BPA and BPA glucuronide concentrations were measured in 10 men by Teeguarden et al. (2015), following administration of 30 µg/kg bw *d*_6_-BPA in 355 mL soup, and previously modeled by Karrer et al. (2018). BPS and BPS glucuronide concentrations were measured after administration of 8.75 μg/kg bw of *d*_4_-BPS in a chocolate cookie, to 4 men and 3 women.^31^ Previous modeling of BPS concentrations is a top-down non compartmental approach. ^31^ Note: BPA, bisphenol A; BPS, bisphenol S; MC, Monte Carlo.

The present model was also evaluated for BPS concentrations predictions, with measures from Oh et al. (2018). Blood C_max_ values predicted by the present model for BPS and its glucuronide were close to observed values: 143 and 94% respectively. Our predicted t_max_ were delayed by 2.5 h (parent compound) and 20 min (metabolite). The predicted AUC for BPS glucuronide closely matched the *in vivo* value, while the predicted AUC for BPS parent was 5 times higher than observed. Three of the six predicted intervals of variability induced by MC simulations overlapped with the measured ranges, notably C_max_ values. These results indicate that the present model generates accurate predictions of BPS concentrations.

### Sensitivity analysis of model parameters

Six parameters exerted significant influence on all bisphenol analogs concentrations in all key organs and for all human physiological models: these were central physiological parameters (body weight and cardiac output) and parameters related to the liver and glucuronidation kinetics (Vmax, Km, liver volume, and liver blood flow) (Figure 6). Other parameters associated with hepatic glucuronidation were frequently sensitive, notably the partition coefficient in the liver of the glucuronide metabolite and the scaling factor of glucuronidation in children (sensitive in all non-adult models). EHC rate was also frequently identified as sensitive. Lastly, parameters associated with organs of toxicological interest (volumes, partition coefficients, and flow rates of thyroid, breasts and testes) were identified as sensitive, but only for the concentrations in the organs to which they are associated.

**Figure 6.**
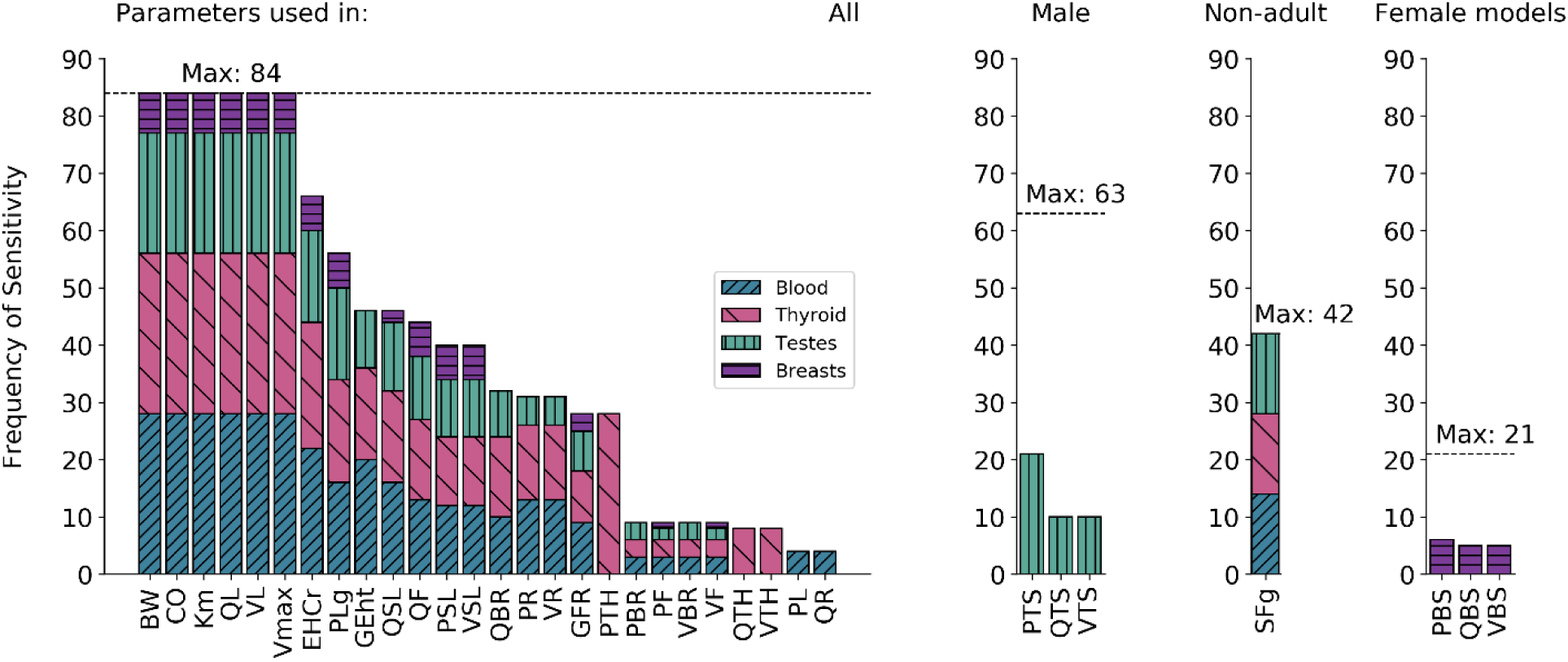
Frequency of sensitivity of parameters across sensitivity analysis in all human models (4) and bisphenols (7) in 4 different compartments (3 organs of interest per physiological model), for exposure scenarios 5 to 8 (Table 1). Dashed horizontal lines indicate the maximum frequency to which the parameters can be found sensitive among the analysis. Note: BW, body weight; CO, cardiac output; EHCr, enterohepatic circulation rate; GEht, gastric emptying half-time; GFR, glomerular filtration rate; KM, Michaelis-Menten constant; PBR, partition coefficient in the brain; PBS, partition coefficient in the breasts; PF, partition coefficient in the adipose tissue; PL, partition coefficient in the liver, PLg, partition coefficient of the glucuronide form in the liver; PR, partition coefficient in the rapidly perfused tissues; PSL, partition coefficient in the slowly perfused tissues; PTH, partition coefficient in the thyroid; PTS, partition coefficient in the testes; QBR, blood debit in the brain; QBS, blood debit to the breasts; QF, blood debit to the adipose tissue; QL, blood debit to the liver; QR, blood debit to the rapidly perfused tissues; QSL, blood debit to the slowly perfused tissues; QTH, blood debit to the thyroid; QTS, blood debit to the testes; SFg, glucuronidation scaling factor; VBR, brain volume; VBS, breasts volume; VF, volume of adipose tissue; VL, liver volume; Vmax, maximum enzyme velocity; VR, volume of the rapidly perfused tissues; VSL, volume of the slowly perfused tissues; VTH, thyroid volume; VTS, testes volume.

### Internal exposure assessments

Concentrations of seven bisphenol analogues were predicted in the blood, thyroid, testes, and breast tissues of man, woman, toddler, and child models following single or repeated dose exposure. Predicted concentrations profiles were similar between age groups, so only man and child model results were represented for simplicity (Figure 7). For all bisphenol analogs, higher concentrations were observed in blood, thyroid and testes in non-adult models compared to adult models (Figure 7, Table 4). Repeated exposure scenarios enabled the observation of bisphenol accumulation in compartment and the attainment of steady state, where the rate of substance intake matched the rate of elimination, resulting in stable tissue concentrations over time (Figure 7, Figure 8, right panel). Steady state was not reached over the course of the 96 h simulation for a majority of bisphenols analogs in breasts (Figure 8), so comparison of the bisphenol analogs accumulation was performed in blood, testes and thyroid.

**Figure 7.**
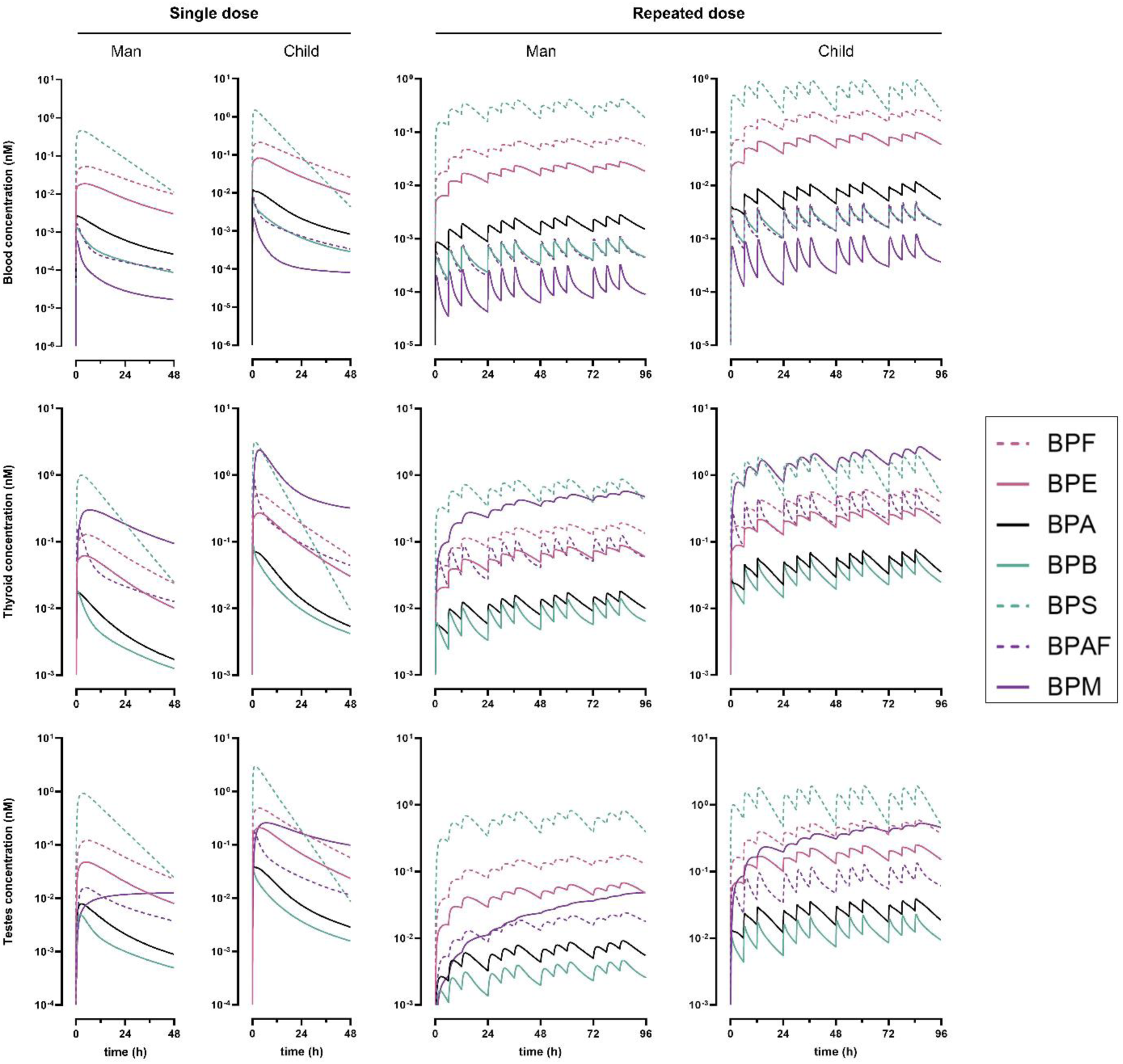
Predicted concentrations profiles (median of 10’000 MC simulations) of BPA, BPAF, BPB, BPE, BPF, BPM and BPS in blood, thyroid and testes in man and child models after single or repeated dose (exposure scenarios 5, 7, 9 and 11 of Table 1). Bisphenols are ordered by ascending MW. Note: BPA, bisphenol A; BPAF, bisphenol AF; BPB, bisphenol B; BPE, bisphenol E; BPF, bisphenol F; BPM, bisphenol M; BPS, bisphenol S; MC, Monte Carlo.

**Figure 8.**
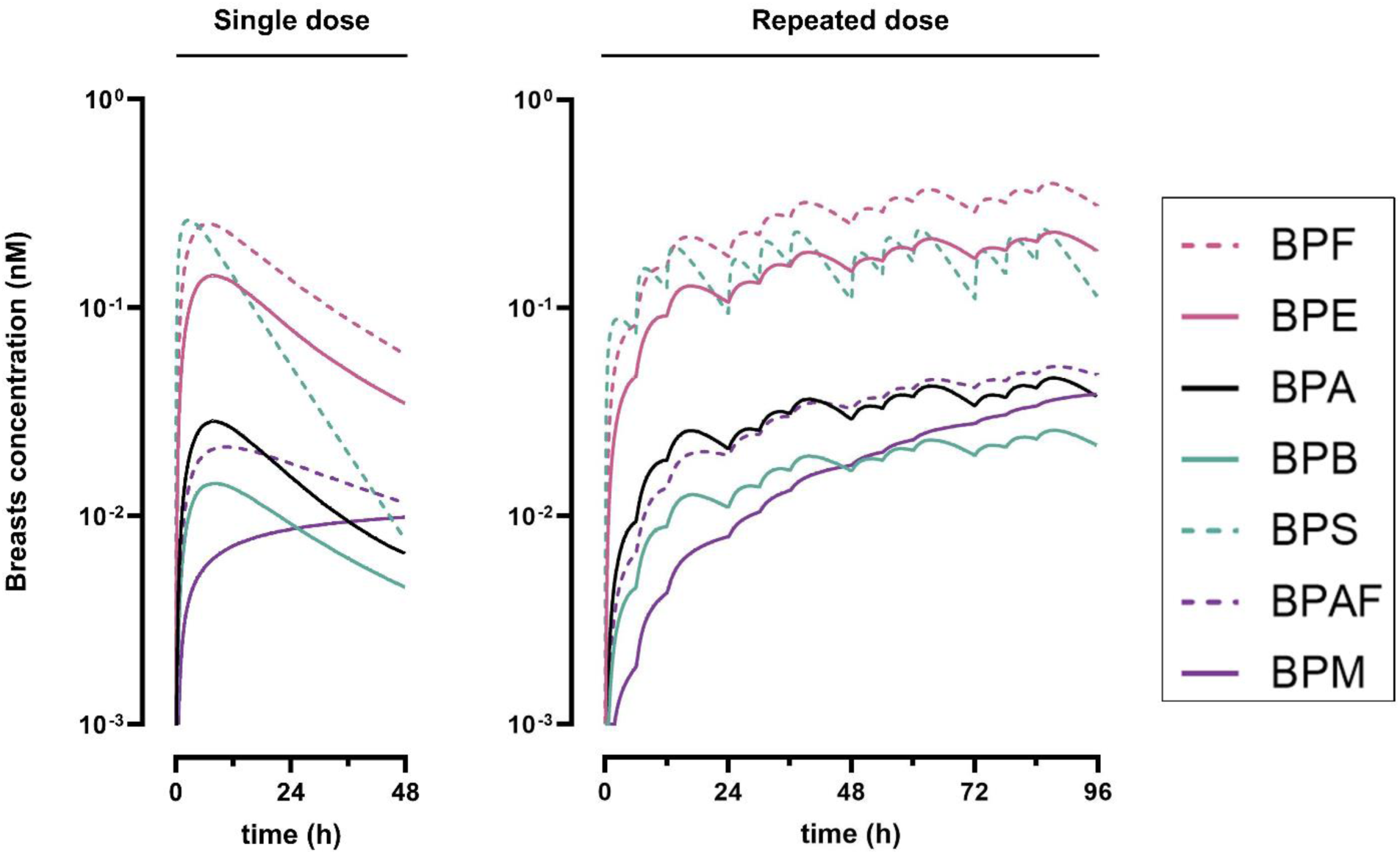
Predicted concentrations profiles (median of 10’000 MC simulations) of BPA, BPAF, BPB, BPE, BPF, BPM and BPS in breasts after single dose exposure after single or repeated dose (exposure scenarios 6 and 10 of Table 1). Bisphenols are ordered by ascending MW. Note: BPA, bisphenol A; BPAF, bisphenol AF; BPB, bisphenol B; BPE, bisphenol E; BPF, bisphenol F; BPM, bisphenol M; BPS, bisphenol S; MC, Monte Carlo.

**Table 4.** Highest C_max_ values per bisphenol and per key organ after repeated dose (exposure scenarios 9 to 12), among the human models. Bisphenols are ordered by ascending MW.

| bisphenol | max. C<sub>max</sub> blood (pM) | max. C<sub>max</sub> breasts (pM) | max. C<sub>max</sub> testes (pM) | max. C<sub>max</sub> thyroid (pM) |
| --- | --- | --- | --- | --- |
| BPS | 938<sup>a,b</sup> | 239<sup>c</sup> | 1921<sup>a,b</sup> | 2017<sup>b</sup> |
| BPF | 260<sup>b</sup> | 397<sup>a,c,d</sup> | 590<sup>b</sup> | 627<sup>b</sup> |
| BPE | 98<sup>b</sup> | 231<sup>c,d</sup> | 251<sup>b</sup> | 318<sup>b</sup> |
| BPA | 11<sup>b</sup> | 46<sup>c,d</sup> | 40<sup>b</sup> | 76<sup>b</sup> |
| BPAF | 5<sup>b</sup> | 52<sup>c,d</sup> | 133<sup>b</sup> | 595<sup>b</sup> |
| BPB | 5<sup>b</sup> | 26<sup>c,d</sup> | 23<sup>b</sup> | 63<sup>b</sup> |
| BPM | 1<sup>e</sup> | 39<sup>c,d</sup> | 531<sup>b,d</sup> | 2665<sup>a,b,d</sup> |

<sup>a</sup> Highest concentration per compartment (among all bisphenols).
<sup>b</sup> Concentration predicted by the child model.
<sup>c</sup> Concentration predicted by the woman model.
<sup>d</sup> Concentration for which the steady state was not reached.
<sup>e</sup> Concentration predicted by the toddler model.
Note: BPA, bisphenol A; BPAF, bisphenol AF; BPB, bisphenol B; BPE, bisphenol E; BPF, bisphenol F; BPM, bisphenol M; BPS, bisphenol S.

Among the seven bisphenol analogues studied, three groups can be identified, each with distinct characteristics in terms of their ranking of concentrations in blood and tissues, as well as the speed at which steady state is reached in compartments. BPS, BPF, and BPE often have the highest concentrations in blood and tissues, reaching steady state the fastest, within 24 to 48 h (Figure 7, Figure 8, Table 4). Predicted concentrations of BPB and BPA are among the lowest in all compartments, especially in thyroid and testes, and reach steady state around 48 h (Figure 7, Figure 8). BPM and BPAF are not exactly similar, but both had predicted concentrations that were the lowest in blood and the highest in thyroid (Figure 7). BPM concentration did not decrease during the 48 h following single dose exposure, and did not reach steady state in thyroid and testes of non-adults model, during the four days of the simulation (Figure 7, Table 4). While the higher concentrations of all other bisphenols were predicted in the child (in blood, testes and thyroid), maximum blood concentration of BPM was predicted in the toddler (Table 4).

## Discussion

In this study, we developed PBK models to predict internal exposure of bisphenols prevalently detected in human matrices: BPA, BPAF, BPB, BPE, BPF, BPM, and BPS. The findings demonstrate that crucial ADME processes, such as intestinal uptake, EHC and glucuronidation kinetics could be parametrized in a multimodal manner that is independent from biomonitoring data and read-across. Moreover, the model evaluation revealed that predicted BPA and BPS concentrations were reliable. Concentrations of bisphenol analogs in key organs (blood, thyroid, testes and breasts) were predicted in man, women, child and toddler, after environmentally relevant oral exposure.

### PBK models parametrization without in vivo data

A variety of methods were employed to parameterize PBK models for a family of compounds, some of which were considered data-poor (Figure 2). QSARs, *in vitro* experiments and cross-species extrapolation were selected for their simplicity of use, requiring minimal prior information (mainly molecular structures). Whenever possible, their results were compared to previously available results to ensure the biological accuracy of our parameters. Additionally, these chosen methods were designed to be robustly applicable to all molecules within the compound class, which should also enhance the reproducibility of this framework for other families of molecules.

The parametrization of intestinal uptake of the bisphenol analogs could be achieved via the estimation of their P_app_ values by various techniques. Predictions of this value by QSARs from Kamiya et al. (2020) and Lanevskij and Didziapetris (2019) were compared together and evaluated against available *in vitro* measures (Figure 3). Predictions were generally consistent between the results of the various techniques, and in line with actual actual transport values measured using Caco-2 monolayers for BPA and BPS. Variability in P_app_ value predicted by the two QSARs (for BPM) and discrepancies between *in silico* predictions and *in vitro* measures (for BPS) stemmed from differing sensitivities of the QSARs to molecular volume and the presence of electronegative atoms, impacting reproducibility of P_app_ prediction. Furthermore, the inclusion of three bisphenols compounds in the training dataset of QSAR K^56^ did not significantly enhance predictions of P_app_ value compared to QSAR L^58^, indicating specialized training sets may not necessarily improve accuracy. Overall, these observations underscore the feasibility of using QSARs for modeling intestinal uptake, highlight the benefits of integrating multiple independent methods and demonstrate the continued value of additional *in vitro* permeability measurements.

To design a model concept that is common for all bisphenols and based on molecule-specific parameters, metabolism parametrization of the present model was based on glucuronidation kinetics measured in hepatic fractions. Bisphenols glucuronidation was assessed either in liver S9 fractions (for BPA, BPB, BPE and BPM), or in liver microsomes (for BPS, BPAF and BPF).^32^ Data generated from those various experiments was considered homogeneous since compositions of homologous UGTs are strongly correlated between commercially available pool human liver microsomes and human liver S9 fractions.^77^ However, BPA glucuronidation rates were faster when catalyzed by human hepatic S9 fractions in the present study. Furthermore, contrary to previous findings, we did not detect that intestinal enzyme preparations catalyze the glucuronidation of BPA, BPB, BPE nor BPM.^32^ Hepatic and intestinal glucuronidation of BPA, BPS, BPAF, and BPF was characterized in human hepatic and intestinal microsomes and cell lines.^32,78–80^ To the best of our knowledge, glucuronidation kinetics for BPB, BPE, and BPM have not been previously reported. The glucuronidation capability of the intestinal S9 system was validated with Urolithin A, which produced Urolithin A-glucuronide.^61^ Thus, intestinal glucuronidation was not included in the model, as it could not be characterized for all bisphenols analogs and previous results indicate that bisphenols are metabolized more in the liver than in the intestine.^32^ Hepatic sulfation was also not represented due to the lack of comprehensive data for all analogs. ^81^ However, the latter have a significant importance in children, particularly in girls,^69^ which is beyond the scope of the present model. Finally, since hepatic glucuronidation parameters were the most sensitive for all model predictions, we confirmed the choice of parametrization using *in vitro* measurements (rather than an *in silico* method) for greater reliability.

In the absence of available tissue-to-blood partition coefficients for all bisphenol analogs in humans,^82^ partitioning was estimated using a uniform approach based on an established QSAR method.^50,52^ Lipophilicity had an important influence on partition coefficient values, similar to the impact observed in the previous PBK model for BPA, BPAF, BPF and BPS,^32^ in which partition coefficient values predicted with the method by Zhang and Zhang (2006). This point was previously suggested as influencing BPAF internal exposures.^32,84^ In the present model, sensitivity analyses results indicated that liver-to-glucuronide metabolite partition coefficient (PLBPg) significantly influences predicted concentrations of original bisphenols, highlighting the need to include compartments that are biologically relevant (i.e. represent actual physiological tissue) for metabolite distribution. Overall, in a data-poor context, determining partition coefficients is largely hindered by the lack of tissue composition data available in literature. Further characterization of sensitive endocrine tissues, such as the ovaries, breasts, and uterus, is needed to enhance the predictive scope of models used in female reproductive toxicity risk assessment.^85,86^

### PBK models evaluation with BPA and BPS biomonitoring

The performance of the present model in predicting blood concentrations of BPA, BPS, and their metabolites was evaluated using human experimental data. Predicted C_max_ had high accuracy for all four molecules (Figure 5). Moreover, the prediction variability obtained by re-sampling sensitive parameters in Monte Carlo simulations, to account for parameter variability and uncertainty, overlapped with the observed ranges of measured values for blood C_max_ (BPA, BPA glucuronide, BPS, BPS glucuronide), t_max_ (BPA, BPA glucuronide), and AUC (BPA, BPA glucuronide, BPS glucuronide). This indicates that the present model provided reliable predictions of BPA and BPS predictions. Furthermore, the present model performances were compared to previous modeling efforts for BPA and BPS. Following oral BPA administration, predictions of toxicokinetic parameters by the present model were closer to the measured values than the predictions of the compartmental model developed by Karrer et al. (2018). Although predictions for BPS toxicokinetics were less accurate than those from the model by Oh et al. (2018), based on experimental data and offering descriptive rather than predictive capabilities, the present model provides a reliable tool for predicting BPS toxicokinetics in humans. Comparing the performance of the present model with previous modeling efforts is valuable for assessing the reliability of predictions from models that use different parameterization approaches. Indeed, comparisons of predicted blood C_max_ values demonstrated that the present multimodal parameterization approach achieves equal or greater similarity to *in vivo* measures than the *in vivo* data fitting methods used in previous models (Figure 5). Moreover, since models built in data-poor situations are not calibrated with the whole available biomonitoring dataset, their predicted internal concentrations can be evaluated against additional internal concentration data, which enhances confidence in their predictions. For instance, the BPS blood concentration values measured by Oh et al. (2018) were used to calibrate the PBK model established by Karrer et al. (2018), and thus, could no longer be used for its evaluation. Thus, our PBK model is the first to be evaluated for two different bisphenols and their metabolites, which ensures more reliable predictions.

Model evaluation results highlighted the need to improve the present model parametrization of ADME processes downstream of systemic distribution. Despite accurate C_max_ predictions, all AUC values were overestimated compared to *in vivo* measures, indicating that urinary excretion and/or EHC were not modeled accurately (Figure 5). Overestimation of EHC is a primary hypothesis to explain the reduced predictive performance of the present model, especially the delayed t_max_. Urinary excretion, parameterized from GFR and adjusted for age and sex, was sensitive in over a quarter of the sensitivity analyses (Figure 6). EHC rate significantly impacted more than two-thirds of the concentrations analyzed (Figure 6), which aligns with its well-established influence on xenobiotic retention within organisms. The motivation to model EHC rate from concentration data measured in rats stemmed from the availability of blood concentration measurements beyond 24 h post-administration and confirmed EHC of bisphenols in these animals.^27^ However, differences in EHC have been observed between rodents and humans. For example, the MW threshold for excreting chemicals via bile is estimated to be 400 Da for rats and 475 Da for humans.^63^ The MW’s of metabolites in this study ranged from 376 to 522 Da (Table S1), with three of the metabolites having MW’s between these thresholds. These interspecies differences may explain the inaccuracies in EHC modeling from rat data. Moreover, while the prolonged elimination time of BPS suggests it undergoes EHC in humans,^32^ this observation was not described for other bisphenols which might not even be substrates for human hepatic canalicular transporters.^87^ Ultimately, the EHC rates used in the present model were higher than previous estimations. In Yang et al. (2015), only ten percent of glucuronidated BPA underwent EHC, a rate seven times lower than our predictions (Table 3). Despite the perfectibility of EHC parametrization, we still considered the present approach to be acceptable since the predicted AUC values were within the range of *in vivo* measurements for three out of four molecules (Figure 5).

### Internal exposure assessments

Kinetic behavior of BPA, BPAF, BPB, BPE, BPF, BPM and BPS were predicted after environmentally relevant oral exposure. Internal concentrations of BPA, BPAF, BPF and BPS were predicted by a different model under similar scenarios of exposures, and our findings align with those previous results.^32^ One particular finding shared by both studies is the prediction of higher internal maximal concentrations in non-adult models, particularly in children and toddlers. Here, this phenomenon in enhanced, due to the application of an age-related scaling factor for UGT2B15 enzyme expression.^69,70^ This slowed down bisphenol glucuronidation, which contributed to the predicted higher concentrations of the original bisphenol in non-adult models.^69,70^

The opposite was observed for predictions of BPA concentrations, which were lower in the present model compared to the previous one, due to more efficient glucuronidation rates.^32^ BPE, BPB, and BPM had not been previously modeled toxicokinetically in humans. Among the seven analogs studied, BPE had the third slowest glucuronidation rate (Figure 4), the second lowest EHC rate (Table 3) and the third lowest partition coefficients, which consequently led to its concentrations being consistently among the highest across most organs (Figures 7 and 8). In contrast, BPB exerted faster glucuronidation rates under physiological conditions and higher blood-tissue partition coefficients (Figure 4). Therefore, it remained longer in the liver, where it was rapidly metabolized and excreted. As a result, BPB concentrations in tissues were among the lowest in all tissues (Figure 7, Figure 8). While BPM underwent rapid hepatic glucuronidation (Figure 4), its kinetics differed due to partitions coefficients that were several degrees of magnitudes higher than other bisphenols analogs, resulting in a marked distinction between its low blood concentration (Figure 7) and high accumulation in other compartments, which persisted beyond 96 h of simulation (Table 4, Figures 7 and 8). Finally, the highest blood concentration of BPM in a repeated dose exposure scenario was predicted in the toddler model, which falls in line with previous observations.^24^

The presented model predicted bisphenol concentrations in three toxicologically relevant compartments related to the toxic properties of BPA: thyroid, breasts (in adult women), and testes (in males). This model is the first to predict concentrations of BPA analogs in the thyroid, despite their thyroid hormone-disrupting potential being previously evaluated *in vitro* at concentrations much higher than those predicted here. For instance, Wu et al. (2022) tested cell viability and thyroid hormone receptors α and β expression levels in HTR-8/SVneo cells with BPA analog concentrations ranging from 1 to 1000 µM. The breasts also represent a compartment where the toxicokinetic properties of BPA alternatives had not been explored, despite the carcinogenic potential of ED in breast tissue. The potential effects of by BPB, BPA, BPF, and BPS on estrogen receptor α, which can lead to tumor progression and metastasis, were studied in cell lines MCF-7, MDA-MB-231, and T47D cells at concentrations ranging from 10^−11^ to 10^−4^ M in the absence of precise exposure data.^89^ Concerningly, BPA was reported to exert increased combinatory estrogenicity in co-exposure with other xenoestrogens.^3,90^ BPA analogues are commonly observed simultaneously in human biomonitoring.^16–18,20–24^ In this light, the current PBK models could serve as a starting point of selecting concentrations of BPA analogues to test for mixture effects and combinatory toxicity.

Although Karrer et al. predicted concentrations of BPA, BPAF, BPF, and BPS in the gonads, our model more precisely parameterized male testes physiology (volume, flow, tissue composition for partition coefficients^53^) and extended this to three other bisphenols of reprotoxicological concern. For instance, the impact of BPA, BPF, BPS, BPE, and BPB on the endocrine function of adult human testis explant cultures was tested by Desdoits-Lethimonier et al. (2017) at concentrations between 10^−9^ and 10^−5^ M. Cytotoxicity and phenotypic marker observations due to BPA, BPAF, BPF, BPS, and BPM exposure in germ and steroidogenic cell lines were also performed at concentrations ranging from 1 nM to 100 µM using single-cell high-content imaging.^92^ Our predictions provide a toxicokinetic perspective to these studies, clarifying their results and support further studies of BPA analogs toxicity.

## Conclusion

The present study advances the toxicokinetic understanding of bisphenol analogs, providing internal concentrations and distribution patterns following oral uptake. The present PBK models demonstrated robust performance in predicting parent compound and glucuronide concentrations, underscoring the feasibility of model parameterization in data-poor situations, particularly through the evaluation of metabolic kinetics, determination of tissue partition coefficients, and incorporation of EHC. In this study, we addressed the toxicokinetics of BPB, BPE, and BPM in humans for the first time. While tissue concentrations of BPB were among the lowest among the bisphenols included in the study, BPE and BPM were predicted to accumulate in high concentration in key organs, especially in kid and toddler models. Finally, predictions of bisphenol concentrations in the thyroid, breasts, and testes offer novel insights for interpreting existing toxicological data, particularly concerning bisphenol-associated endocrine disruption and associated carcinogenic potential. By providing a detailed toxicokinetic perspective, the models created in this study are expected to contribute to a more precise evaluation of health risks induced by bisphenol analogs, guiding their safer use.

