## Supplementary Material for "Human internal exposures of bisphenol A and six data-poor analogues predicted by physiologically based kinetic models with multimodal parameterization"

### Figure S1: Workflow of PBK model development, validation, reporting and dissemination (adapted from OECD (2021)).

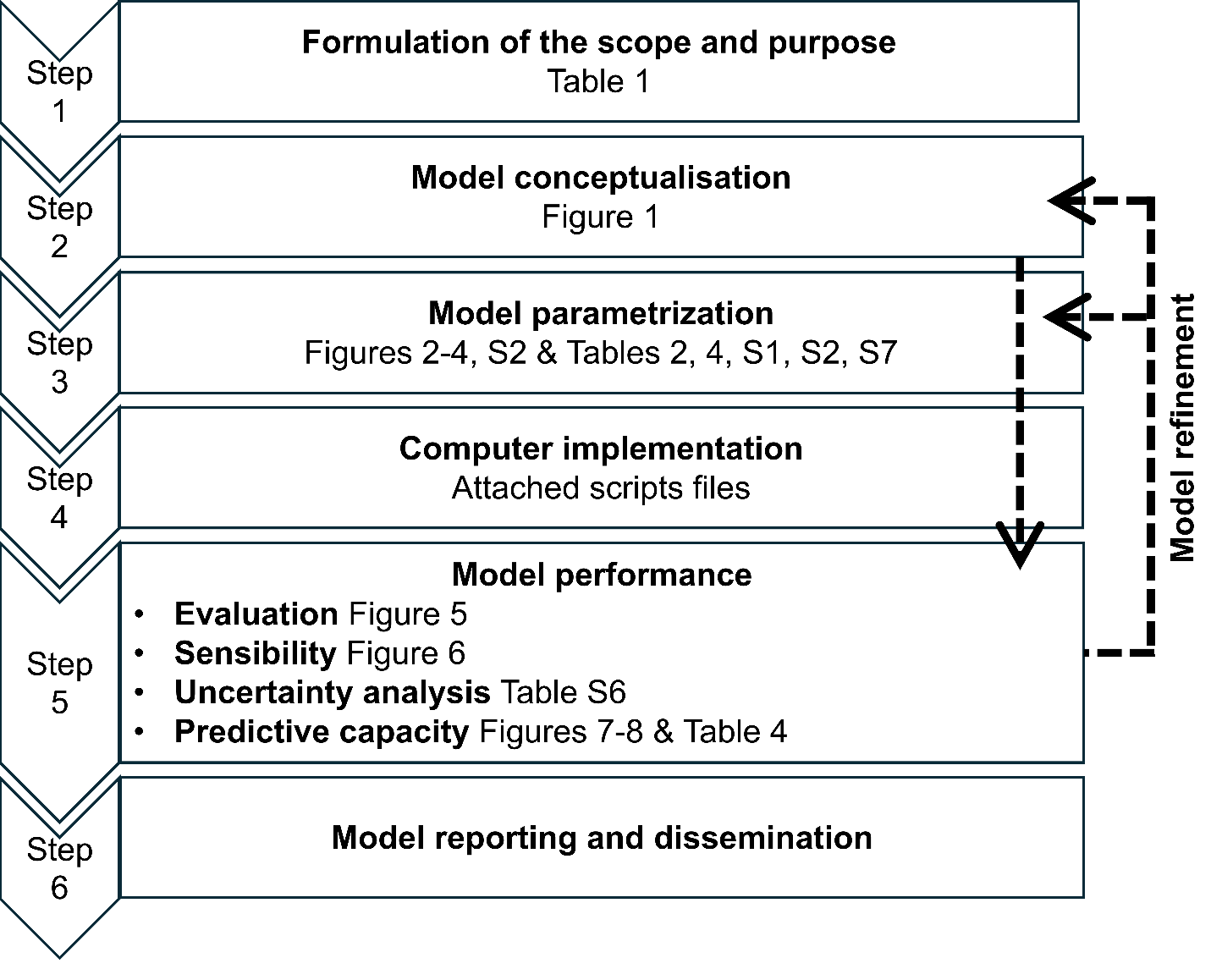

### Table S1: Physicochemical properties of bisphenol glucuronides (predicted by ChemDraw 20.0).

|  | **MW (g/mol)** | **pKa** | **logP** |
| --- | --- | --- | --- |
| **BPA glucuronide** | 404.42 | 9.509 | 2.27 |
| **BPAF glucuronide** | 512.36 | 8.897 | 3.13 |
| **BPB glucuronide** | 418.44 | 9.487 | 2.68 |
| **BPE glucuronide** | 390.39 | 9.58 | 1.79 |
| **BPF glucuronide** | 376.36 | 9.651 | 1.46 |
| **BPM glucuronide** | 522.59 | 9.474 | 5.16 |
| **BPS glucuronide** | 426.39 | 8.429 | 0.26 |

### Table S2: Physiological parameters.

|  | MAN | WOMAN | CHILD | TODDLER | RAT |
| --- | --- | --- | --- | --- | --- |
| - ***Fractional organ volumes (convert human to %)*** | | | | | |
| VLC - Liver volume | 0.0319 | 0.0298 | 0.0374 | 0.0380 | 3.4 |
| VFC - Adipose volume | 0.2036 | 0.3055 | 0.1932 | 0.2343 | 7 |
| VSKC - Skin volume | 0.0474 | 0.0403 | 0.0322 | 0.0365 | 19.7 |
| VBRC - Brain volume | 0.0207 | 0.0224 | 0.0698 | 0.0956 | 0.6 |
| VSTC- Stomach volume | 0.0028 | 0.0030 | 0.0034 | 0.0025 | 0.5 |
| VTSC- Testes volume | 0.0005 | - | 0.0001 | 0.0002 | - |
| VBSC - Breasts volume | - | 0.0083 | - | - | - |
| VTHC - Thyroid volume | 0.0003 | 0.0003 | 0.0002 | 0.0002 | - |
| VBC - Blood volume | 0.0726 | 0.0650 | 0.0737 | 0.0500 | 7.4 |
| VRC - Rapidly perfused tissues volume | 0.0269 | 0.0258 | 0.0283 | 0.0278 | 1.5 |
| VSLC - Slowly perfused tissues volume | 0.5563 | 0.4330 | 0.4381 | 0.3158 | 47.7 |
| VGLC - Gut lumen volume | 0.0089 | 0.0100 | 0.0114 | 0.0173 | 0.9 |
| VGTC - Gut tissue volume | 0.0140 | 0.0160 | 0.0179 | 0.0135 | 1.6 |
| - ***Intestinal surface areas (SA) (dm2)*** | | | | | |
| SASI small intestine | 2917.5 | 2709.1 | 1771.3 | 1250.3 | - |
| SALI large intestine | 107.8 | 107.8 | 73.5 | 58.8 | - |
| ISA - total intestinal SA (dm2) | 3025.3 | 2816.9 | 1844.8 | 1309.1 | 1.44 |
| - ***Further physiological parameters*** | | | | | |
| TRANSI - transit time (h) | 40 | 52 | 38 | 36 | 18 |
| GEst - time to empty 50% of stomach (min) | 11.8 | a | a | a | 25 |
| GFR (ml/min [/1.73 m2 for human]) | 125 | a | a | a | 1.31 |
| CO - Cardiac output (L/min) | 6.5 | 5.9 | 3.4 | 1.2 | 0.083 |
| - ***Fractional blood flows (% or cardiac output)*** | | | | | |
| QLC - Liver debit | 6.5 | 6.5 | b | b | 18.3 |
| QFC - Adipose debit | 5 | 8.5 | b | b | 7.0 |
| QRC - Rapidly perfused tissues debit | 123 | 122 | b | b | 21.3 |
| QSLC - Slowly perfused tissues debit | 22 | 17 | b | b | 40 |
| QSKC - Skin debit | 5 | 5 | b | b | 5.8 |
| QGTC - Gut tissue debit | 14 | 16 | b | b | 9.0 |
| QBRC - Brain debit | 12 | 12 | b | b | 2.0 |
| QTHC - Thyroid debit | 1.5 | 1.5 | b | b | - |
| QTSC - Testes debit | 0.05 | - | b | b | - |
| QBSC - Breast debit | - | 0.4 | - | - | - |

a: Value conserved for all human models

b: Value conserved for all male models

### Table S3: Partition coefficients (P-) and unbound fractions (fu, FU-).

|  | **BPS** | **BPF** | **BPE** | **BPA** | **BPB** | **BPAF** | **BPM** |
| --- | --- | --- | --- | --- | --- | --- | --- |
| Adipose tissue (PFBP) | 2.04 | 17.76 | 30.13 | 63.30 | 122.17 | 235.07 | 5537.21 |
| Brain (PBRBP) | 1.47 | 7.69 | 11.91 | 22.04 | 38.03 | 66.41 | 895.40 |
| Breasts (PBSBP) | 0.50 | 4.35 | 7.38 | 15.50 | 29.93 | 57.59 | 1356.62 |
| Gut tissue (PGTBP) | 1.60 | 8.06 | 12.43 | 22.92 | 39.47 | 68.86 | 927.02 |
| Liver (parent, PLBP) | 0.87 | 4.06 | 6.24 | 11.49 | 19.76 | 34.46 | 463.79 |
| Liver (glucuronide, PLBPgluc) | 0.49 | 0.55 | 0.67 | 1.06 | 1.72 | 2.98 | 42.17 |
| Rapidly perfused (PRBP) | 0.97 | 4.41 | 6.74 | 12.34 | 21.18 | 36.86 | 495.08 |
| *Lungs* | *1.08* | *4.98* | *7.62* | *13.96* | *23.95* | *41.69* | *559.89* |
| *Kidneys* | *0.87* | *3.83* | *5.85* | *10.72* | *18.41* | *32.06* | *430.83* |
| *Heart* | *0.77* | *3.32* | *5.06* | *9.24* | *15.84* | *27.56* | *369.79* |
| Skin (PSKBP) | 2.31 | 12.03 | 18.55 | 34.23 | 58.93 | 102.79 | 1383.46 |
| Slowly perfused (PSLBP) | 0.64 | 2.72 | 4.16 | 7.61 | 13.09 | 22.81 | 306.74 |
| *Muscles* | *0.60* | *2.43* | *3.7* | *6.77* | *11.63* | *20.26* | *272.48* |
| *Bones* | *0.74* | *3.52* | *5.4* | *9.93* | *17.08* | *29.76* | *400.31* |
| Testes (PTSBP) | 2.17 | 2.30 | 2.56 | 3.44 | 5.47 | 33.55 | 870.31 |
| Thyroid (PTHBP) | 2.28 | 2.45 | 3.30 | 6.58 | 14.46 | 132.39 | 3655.37 |
| fu (parent, FUBP) | 0.25 | 0.09 | 0.06 | 0.04 | 0.03 | 0.02 | 0.002 |
| fu (glucuronide, FUBPG) | 0.70 | 0.40 | 0.32 | 0.22 | 0.16 | 0.11 | 0.01 |

### Table S4: Chemicals, Reagents and Enzymes used for glucuronidation kinetics

| **Material** | **Supplier** |
| --- | --- |
| Corning UltraPool Human Liver S9 (mixed gender, 150-donorpool) | Corning |
| Corning Gentest UGT Reaction Mix | Corning |
| Pooled human intestinal S9 fraction | Biopredic |
| Uridine-5’-diphosphoglucuronic acid (UDPGA) trisodium salt | Sigma–Aldrich |
| Magnesiumchloride | Sigma–Aldrich |
| ß-Glucuronidases (from bovine liver) | Sigma–Aldrich |
| Sodium acetate | Sigma–Aldrich |
| Alamethicin | Enzo Life Sciences AG |
| Urolithin A (3,8-Dihydroxy-6H-benzo[c]chromen-6-one) | abcr |
| Bisphenol A | Sigma–Aldrich |
| Bisphenol AF | Sigma–Aldrich |
| Bisphenol B | Sigma–Aldrich |
| Bisphenol E | Sigma–Aldrich |
| Bisphenol F | Sigma–Aldrich |
| Bisphenol M | Sigma–Aldrich |
| Bisphenol S | Sigma–Aldrich |
| DMSO | VWR |
| Acetonitrile (ACN, HPLC-grade) | Merck-Millipore |
| Tris–HCl | Fluka Chemikals |
| Rat liver S9 (Sprague-Dawley, male) | Sigma-Aldrich |
| Formic acid | Fischer scientific |

### Table S5. HPLC retention times of bisphenols.

| **Bisphenol** | **Retention time tr (min)** | |
| --- | --- | --- |
|  | **Parent** | **Metabolite** |
| BPA | 14.8 | 10.1 |
| BPAF | 18.1 | 13.1 |
| BPB | 16.6 | 11.4 |
| BPE | 13.5 | 9.1 |
| BPF | 12.0 | 7.8 |
| BPM | 22.4 | 17 |

### Table S6: Variability of sensible parameters.

| **Parameter** | | **CV (%)** | **Distribution** | **Reference** |
| --- | --- | --- | --- | --- |
| BW | Body weight (kg) | 26 | Lognormal | (Clewell et al., 1999) |
| - Organ volumes (fraction of BW) | | | | |
| VL | Liver volume | 25 | Normal | (Clewell et al., 1999) |
| VR | Rapidly perfused tissues volume | 30 | Normal | (Clewell & Clewell, 2008) |
| VSL | Slowly perfused tissues volume | 16 | Normal | (Clewell et al., 1999) |
| VTS | Testes volume | 30 | Normal | (Clewell & Clewell, 2008) |
| VBR | Brain volume | 30 | Normal | (Clewell & Clewell, 2008) |
| VF | Adipose volume | 24 | Normal | (Clewell et al., 1999) |
| VTH | Thyroid volume | 30 | Normal | (Clewell & Clewell, 2008) |
| VBS | Breasts volume | 30 | Normal | (Clewell & Clewell, 2008) |
| - Further physiological parameters | | | | |
| CO | Cardiac output (L/min) | 22 | Normal | (Clewell et al., 1999) |
| EHCr | EHC rate | 0.59 | Lognormal | (Guiastrennec et al., 2018) |
| GEst | Time to empty 50% of stomach (min) | 69 | Lognormal | (Oberle et al., 1990) |
| GFR | GFR (ml/min [/1.73 m2 for human]) | 18 | Lognormal | (Fravel et al., 2023) |
| - Fractional blood flows (% or cardiac output) | | | | |
| QL | Liver debit | 32 | Normal | (Clewell et al., 1999) |
| QSL | Slowly perfused tissue debit | 30 | Normal | (Clewell & Clewell, 2008) |
| QF | Adipose debit | 30 | Normal | (Clewell & Clewell, 2008) |
| QBR | Brain debit | 30 | Normal | (Clewell & Clewell, 2008) |
| QTS | Testes debit | 30 | Normal | (Clewell & Clewell, 2008) |
| QTH | Thyroid debit | 30 | Normal | (Clewell & Clewell, 2008) |
| QBS | Breasts debit | 30 | Normal | (Clewell & Clewell, 2008) |
| QR | Rapidly perfused tissues debit | 30 | Normal | (Clewell & Clewell, 2008) |
| - Partition coefficients (tissue-to-plasma) | | | | |
| PLg | Partition of glucuronide to liver | 30 | Lognormal | (Clewell et al., 1999) |
| PSL | Partition of parent BP to slowly perfused tissue | 30 | Lognormal | (Clewell et al., 1999) |
| PR | Partition of parent BP to rapidly perfused tissue | 30 | Lognormal | (Clewell et al., 1999) |
| PTH | Partition of parent BP to thyroid | 20 | Lognormal | (Clewell & Clewell, 2008) |
| PTS | Partition of parent BP to testes | 20 | Lognormal | (Clewell & Clewell, 2008) |
| PF | Partition of parent BP to adipose | 30 | Lognormal | (Clewell et al., 1999) |
| PBR | Partition of parent BP to brain | 30 | Lognormal | (Clewell et al., 1999) |
| PBS | Partition of parent BP to breasts | 20 | Lognormal | (Clewell & Clewell, 2008) |
| PL | Partition of parent BP to liver | 30 | Lognormal | (Clewell et al., 1999) |
| - Glucuronidation kinetics | | | | |
| Vmax | Maximum velocity (nmol/min/mg) | 50 | Lognormal | (Thomas et al., 1996) |
| Km | Michaelis constant (nM) | 20 | Lognormal | (Thomas et al., 1996) |
| SFg | Scaling factor for age | 69 | Lognormal | (Bhatt et al., 2019) |

### Figure S2: Hepatic glucuronidation kinetics.

BPAF and BPF were measured in rat liver S9 fractions, BPM, BPE, BPA and BPB were measured in human liver S9 fractions.

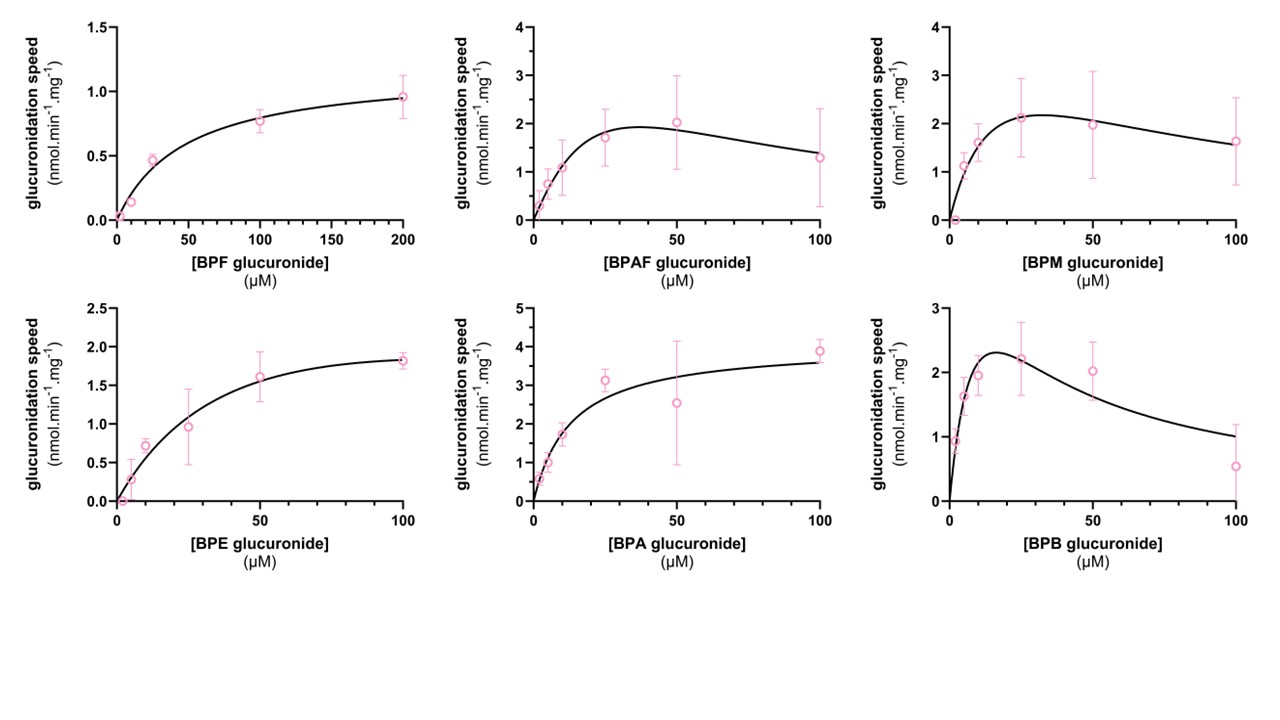

### Table S7: Kinetics parameters for hepatic glucuronidation (fitting from GraphPad Prism).

BPF and BPF values were measured in rat liver S9 fractions and used in rat models. BPA, BPB, BPE and BPM were measured in human liver S9 fractions and used in human models. MM: Michaelis Menten, SI: substrate inhibition.

|  | **BPA** | **BPB** | **BPE** | **BPM** | **BPF** | **BPAF** |
| --- | --- | --- | --- | --- | --- | --- |
| Fitted curve | MM | SI | SI | SI | MM | SI |
| R2 | 0.90 | 0.81 | 0.98 | 0.92 | 0.99 | 0.97 |
| Vmax (nmol.min^-1^.mg^-1^) | 4.06 | 5.81 | 3.42 | 4.89 | 1.17 | 7.10 |
| Km (µM) | 13.10 | 12.34 | 51.31 | 20.15 | 47.54 | 49.68 |
| Ksi (µM) | - | 21.48 | 280.6 | 51.75 | - | 27.63 |

### Model code (Berkeley Madonna)

; PBK model for bisphenols (here BPF) in rat, modeling exposure described in (Lee et al., 2022)

```
; Species: Rat
; Molecule: BPF
; Date: 2024.02.19
; Compiled by Hélène Bigonne

{Top model}
   {Reservoirs}

; ST: stomach compartment
   d/dt (ASTBP) = - ST_outtake
      INIT ASTBP = A0DOSEBP
      LIMIT ASTBP >= 0

; GL: gut lumen compartment
   d/dt (AGLBP) =  ST_outtake - GL_outtake + EHC -FEC
      INIT AGLBP = 0
      LIMIT AGLBP >= 0

; GT: gut tissue compartment
   d/dt (AGTBP) =  GL_outtake - GT_outtake
      INIT AGTBP = 0
      LIMIT AGTBP >= 0

; L: liver compartment
   d/dt (ALBP) =  GT_outtake - L_BPgluc - venL + artL
      INIT ALBP = 0
      LIMIT ALBP >= 0

; B: blood compartment
   d/dt (ABBP) = + venSK + venF + venBR + venL + venSL + venR - artSK - artR - UR_parent - artL - artF - artBR - artSL
      INIT ABBP = 0
      LIMIT ABBP >= 0

; BR: brain compartment
   d/dt (ABRBP) = + artBR - venBR
      INIT ABRBP = 0
      LIMIT ABRBP >= 0

; SL: slowly perfused tissues compartment
   d/dt (ASLBP) = + artSL - venSL
      INIT ASLBP = 0
      LIMIT ASLBP >=0
 ; R: rapidly perfused tissues compartment
   d/dt (ARBP) = - venR + artR
      INIT ARBP = 0
      LIMIT ARBP >= 0

; F: adipose tissue compartment
   d/dt (AFBP) = + artF - venF
      INIT AFBP = 0
      LIMIT AFBP >= 0

; SK: skin compartment
  d/dt (ASKBP) = - venSK + artSK
      INIT ASKBP = 0
      LIMIT ASKBP >= 0

; FECES: fecal compartment
  d/dt (AFECES) = + FEC
      INIT AFECES = 0
      LIMIT AFECES >= 0

; URINE_parent: urine compartment
   d/dt (AURINE_parent) = + UR_parent
      INIT AURINE_parent = 0
      LIMIT AURINE_parent >= 0

   {Flows}

; gastro_intestinal tract flows
   ST_outtake = kelST *ASTBP
   GL_outtake = ka * CVSIBP
   GT_outtake = QGT *CVGTBP

; blood flows
   venL = QL*CVLBP
   artL = QL*CBBP
   artSL = QSL*CBBP
   artBR = QBR*CBBP
   artR = QR*CBBP
   artF = QF*CBBP
   venBR = QBR*CVBRBP
   venSL = QSL*CVSLBP
   venR = QR*CVRBP
   venF = QF*CVFBP
   venSK = QSK*CVSKBP
   artSK = QSK*CBBP

; excretion flows
   UR_parent = CBBP*urineBP
   FEC = AGLBP*KSItransit

   {Submodel "S1"}
      {Reservoirs}

; LBPgluc: liver compartment for BP glucuronide
d/dt (ALBPgluc) = + L_BPgluc - venL_gluc -BILE_intake
    INIT ALBPgluc = 0
    LIMIT ALBPgluc >= 0

; BILE: bile compartment
d/dt (ABILEBPgluc) = + BILE_intake -EHC
    INIT ABILEBPgluc = 0
    LIMIT ABILEBPgluc >= 0

; BBPgluc: blood compartment for BP glucuronide
d/dt (ABBPgluc) = - UR_gluc + venL_gluc
    INIT ABBPgluc = 0
    LIMIT ABBPgluc >= 0

; UrineBPgluc: urine compartment for BP glucuronide
d/dt (AUrineBPgluc) = + UR_gluc
    INIT AUrineBPgluc = 0
    LIMIT AUrineBPgluc >= 0

      {Flow}
BILE_intake = EHCr*CVLBPG*QL 	; biliary excretion
EHC = ABILEBPgluc			; EHC flow from bile
venL_gluc = (1-EHCr)*CVLBPG*QL
L_BPgluc = VmaxLBPFGluc* CVLBP/(Km + CVLBP)
UR_gluc = urineBP*CBBPG


{Globals}
;=====================================================================
; Physiological parameters (rat)
;=====================================================================
```

BW = 0.25	; bodyweight (kg) (Brown et al., 1997)

GutSA= 144	; gut surface area at age 11 weeks (cm2) (Meshkinpour et al., 1981)

;---------------------------------------------------------------------

; relative tissue volumes

;---(fraction of BW)

VLc = 0.034	;  liver (Brown et al., 1997)

VRc = 0.0151	;  rapidly perfused tissues (heart, kidneys, lungs) (Brown et al., 1997)

VSLc = 0.477	;  slowly perfused tissue (bone, muscle) (Brown et al., 1997)

VFc = 0.07	;  fat tissue (adipose) (Brown et al., 1997)

VPc = 0.074	;  blood (Brown et al., 1997)

VSKc = 0.197	;  skin (Brown et al., 1997)

VBRc = 0.006	;  brain (Brown et al., 1997)

VSTc = 0.0052	; stomach (Oatley & Toates, 1969)

VGTc = 0.0162	; gut tissue (Oatley & Toates, 1969)

;---(mL)

VGLc = 2.18	; (mL) Vfluid SI lumen (Tanaka et al., 2020)

```
;---------------------------------------------------------------------
; calculated tissue volumes (L or Kg)
VL = VLc*BW	; liver
VP = VPc*BW	; plasma
VR = VRc*BW	; rapidly perfused tissue
VSL = VSLc*BW	; slowly perfused tissue
VF = VFc*BW	; fat tissue
VSK= VSKc*BW	; skin
VBR = VBRc*BW	; brain
VST = VSTc*BW	; stomach
VGT = VGTc*BW 	; gut tissue
VGL= VGLc/1000	; gut lumen

;---------------------------------------------------------------------
; blood flow rates
```

QC = 0.235*BW^(0.75)*60 ; cardiac output (L/h) (Brown et al., 1997)

;--(fraction of QC)(Brown et al., 1997)

```
QLc = 0.183	; liver
QFc = 0.07		; fat
QRc = 0.213	; rapidly perfused tissue (heart, kidneys, lungs)
QSLc = 0.4		; slowly perfused tissues (bone, muscle)
QSKc = 0.058	; skin
QBRc = 0.02	; brain
;-- (ml/min)
```

QGTc = 7.5	; (ml/min) gut (Davies & Morris, 1993)

```
;---------------------------------------------------------------------
; calculated blood flows (L/h)
QL = QLc*QC		; liver
QF = QFc*QC		; fat tissue
QR = QRc*QC 		; rapidly perfused
QSL = QSLc*QC 		; slowly perfused
QSK = QSKc*QC		; skin
QBR = QBRc*QC		; brain
QGT = QGTc/1000*60	; gut

;=====================================================================
; Physicochemical parameters (BPF)
;=====================================================================

; molecular weights (g/mol)
MWBP = 200.24		; parent compound
MWBPGluc = 376.36	; glucuronidated metabolite

; pKa
pKaBP = 9.658		; parent compound
pKaBPGluc = 9.651	; glucuronidated metabolite

; logP
logPBP = 3.35		; parent compound
logPBPGluc = 1.46	; glucuronidated metabolite

; logD (parent compound)
logDapical = 3.61
logDbasal = 3.6
```

; partition coefficients (parent compound) (Punt et al., 2021; Rodgers & Rowland, 2006)

```
PLBP = 4.06	; liver/blood
PFBP = 17.76	; fat/blood
PRBP = 4.41	; rapidly perfused tissue/blood
PSLBP = 2.72	; slowly perfused tissue/blood
PBRBP = 7.69	; brain/blood
PSKBP = 12.03	; skin/blood
PGTBP = 8.06	; gut/blood
```

; partition coefficients (metabolite) (Punt et al., 2021; Rodgers & Rowland, 2006)

PLBPG = 0.55	; liver/blood

; unbound fractions (Lobell & Sivarajah, 2003; Punt et al., 2021)

```
FUPBP = 0.086	; parent compound
FUPBPG = 0.4	; metabolite
;=====================================================================
; Kinetic parameters
;=====================================================================
; metabolic parameters
; scaling factors
L=VLc*1000	; gram liver /kg BW
VLS9 = 165 	; mg S9/ g liver
; hepatic glucuronidation
Km = 47540	; nM
Vmax = 1.173 ;nmol/min/mg
VmaxLBPFGluc = Vmax*VLS9*60*L*BW

;---------------------------------------------------------------------
; absorption/transfer rates

; intestinal intake (Ka)
```

Papp= 10^(3-0.0038*MWBP+0.41*logDapical-0.3*logDbasal)/10^(7)	;  (cm/s) (Kamiya et al., 2020)

Peff=10^(0.4926*LOG10(Papp) -0.1454)	; (cm/s) (Sun et al., 2002)

```
Ka = Peff*GutSA/1000*360 				; (L/h)

; gastric emptying (kelst)
```

GEstc = 25	; (min) half time of meal in rat stomach (Purdon & Bass, 1973)

```
GEst = GEstc /60		; (h)
LNhalf= -0.69314718		; LN(1/2)
kelST =  -LNhalf/ GEst 	; (h-1) constant of gastric excretion
; from Ct=C0*e^(-kel*t)

; intestinal transit (KSItransit)
```

tpassgut = 18	; (h) passage time through SI+LI (DeSesso & Jacobson, 2001)

```
KSItransit = 1/tpassgut		; (h-1)

;--------------------------------------------------------------------
; EHC parameters
EHCr = 0			; fraction of BPFG formed subject to EHC
;---------------------------------------------------------------------
; urinary excretion
```

GFR = 1.31	;ml/min (Davies & Morris, 1993)

```
urineBP = GFR*(60/1000)	;L/h

;=====================================================================
```

; Run settings (Lee et al., 2022)

```
;=====================================================================
; oral dose
DOSEBPc = 200000000		;ng/kg BW
DOSEBP = DOSEBPc*BW		;ng
A0DOSEBP = DOSEBP/MWBP	;nmol

;====================================================================
; Main model calculations/dynamics: BP
;=====================================================================
; stomach compartment
CSTBP = ASTBP/VST
;---------------------------------------------------------------------
; gut lumen compartment
CGLBP = AGLBP/VGL
;---------------------------------------------------------------------
; gut tissue compartment
CGTBP = AGTBP/VGT		; nmol/L ; BPF concentration in gut
CVGTBP = CGTBP/PGTBP		; partition with gut tissue
;--------------------------------------------------------------------
; liver compartment
CLBP = ALBP/VL
CVLBP = CLBP/PLBP
;---------------------------------------------------------------------
; fat compartment
CFBP = AFBP/VF
CVFBP = CFBP/PFBP
;--------------------------------------------------------------------
; skin compartment
CSKBP = ASKBP/VSK
CVSKBP = CSKBP/PSKBP
;---------------------------------------------------------------------
; rapidly perfused tissue
CRBP = ARBP/VR
CVRBP = CRBP/PRBP
;--------------------------------------------------------------------
; slowly perfused tissue
CSLBP = ASLBP/VSL
CVSLBP = CSLBP/PSLBP
;---------------------------------------------------------------------
; brain
CBRBP = ABRBP/VBR
CVBRBP = CBRBP/PBRBP
;---------------------------------------------------------------------
; blood compartment
CBBP=ABBP/VP			; nmol/L (total)
CBBP2=CBBP*FUPBP			; nmol/L (unbound)
CB1=(CBBP*MWBP)/1000		; ng/mL (total)
CB2=CB1*FUPBP			; ng/mL (unbound)

;=====================================================================
; Sub-model calculations/dynamics: BPgluc
;=====================================================================
; liver
CLBPG = ALBPgluc/VL			;nmol/L
CVLBPG = CLBPG/PLBPG
; blood
CBBPG = ABBPgluc/VP  			;nmol/L
;=====================================================================

{End Globals}
```
